# Enhanced liver regeneration via targeted mRNA delivery for partial in vivo reprogramming

**DOI:** 10.1101/2025.11.03.686120

**Authors:** Beom-Ki Jo, Young Seok Song, Woohyun Song, Hee-Ji Eom, Yon Jae Lee, Jumee Kim, Seunghee Hong, Seung-Woo Cho, Hyuk-Jin Cha

**Author notes:** These authors contributed equally to this work (Beom-Ki Jo, Young Seok Song, Woohyun Song). To whom correspondence should be addressed to Prof. Hyuk-Jin Cha, Ph.D. College of Pharmacy, Seoul National University 1 Gwanak-ro, Gwanak-gu, Seoul 08826, Republic of Korea; Prof. Seunghee Hong, Ph.D. Department of Biochemistry, College of Life Science and Biotechnology, Yonsei University 50, Yonsei-ro, Seodaemun-gu, Seoul 03722, Republic of Korea Prof. Seung-Woo Cho, Ph.D. Department of Biotechnology, College of Life Science and Biotechnology, Yonsei University 50 Yonsei-ro, Seodaemun-gu, Seoul 03722, Republic of Korea.

## Abstract

Recent studies suggest that injury-induced dedifferentiation, which leads to the formation of ‘injury-responsive cells’, contributes significantly to tissue repair across various organs, including the liver. Utilizing Yamanaka factors (Oct4, Sox2, Klf4, and c-Myc: OSKM) for in vivo partial reprogramming generates ‘injury-responsive cells’ in the intestine, mirroring those derived from injury-induced dedifferentiation. Thus, the transgene induction of OSKM or viral delivery of Oct4, Sox2, and Klf4 shows promise in facilitating tissue regeneration in the intestine, liver, skeletal muscle, and retina. Herein, we demonstrated that transient OSKM induction produces two distinct liver progenitor-like cell populations. One of these populations resembles liver progenitor-like cells (LPLCs) generated by acute acetaminophen (APAP) injury without triggering immune responses. To explore in vivo reprogramming as a viable strategy for tissue regeneration, we employed lipid nanoparticles (LNP) carrying OSKM mRNA (OSKM mRNA-LNP) to stimulate LPLCs formation. Notably, the production of Sox9+ LPLCs, and OSKM-induced dedifferentiation, was closely correlated with successful tissue regeneration in the liver post APAP injury. Thus, the OSKM mRNA-LNP approach represents a promising therapeutic intervention for the repair of acute liver injuries.

## Introduction

Mammals are generally acknowledged to exhibit limited tissue regeneration compared to amphibians, which can undergo extensive cellular reprogramming to generate pluripotent-like stem cells, known as blastema, following serious injuries (e.g., amputation)^1^. Nevertheless, recent advancements in single-cell analysis have identified a novel cell population with regeneration potential, termed ‘injury-responsive cells’, in mouse models following acute injuries. This adaptive cellular reprogramming^2^ promotes dedifferentiation, resulting in the formation of injury-responsive cells with fetal-like gene signatures in various organs, including the intestine (revival stem cells: revSCs)^3^, lung (damage associated transient progenitors: DATPs)^4^, and liver (liver progenitor-like cells: LPLCs)^5^. Notably, cytokines from adjacent active immune cells, including IL-6 from Kupffer cells^5^, IL-1β from macrophages, and TGFβ from monocytes or macrophages^6^, play critical roles in triggering dedifferentiation and inducing transient plasticity in the liver, lung, and intestine post-injury, respectively. Similarly, an increase in prostaglandin E2 (PGE2) from rare pericryptal fibroblasts following intestinal injury^7^ drives the production of injury-responsive cells for intestinal regeneration^8^.

Extensive research has been conducted on in vivo cellular reprogramming via the transient induction of Yamanaka factors (Oct4, Sox2, Klf4, and c-Myc: OSKM, or OSK without c-Myc) for tissue regeneration across various organs, including the retina^9^, skeletal muscle^10^, heart^11^, liver^12^, and intestine^13^, as well as for rejuvenation^14^. A recent study revealed that in vivo reprogramming leads to the generation of fetal-like cells, reminiscent of ‘injury-responsive cells’ observed in the injured intestine due to autonomous PGE2 signalling from the intestinal epithelium^13^. This contrasts with post-injury intestinal regeneration, which is primarily driven by the paracrine effect of PGE2 from pericryptal fibroblasts^7^. These findings suggest a shared molecular mechanism underlying dedifferentiation induction after injury and forced cellular reprogramming.

The liver, unlike many mammalian tissues with limited regenerative capacities, is renowned for its remarkable regenerative potential following various injuries^15^. While dedicated stem cells in actively renewing organs, such as the intestine, bone marrow and skin, continuously support fast physiological turnover, the liver, with a slow turnover in adulthood, exhibits marginal renewal effects from stem cells during normal homeostasis, with only a few exceptions in reported literature^16,17^. Following an acute injury, not only hepatocytes proliferation^18^ but also the emergence of LPLCs, which are involved in facultative regeneration in the liver, has been extensively investigated^19^. Genetic and lineage tracing mouse models have demonstrated that LPLCs acting as injury-responsive facultative stem cells^20^ originate from the biliary epithelial cells (i.e., cholangiocytes)^21^ or hepatocytes^22^. Subsequent studies have identified specific types of hepatocytes (e.g., hybrid periportal hepatocytes^23^ and hepatocytes with high telomerase expression^24^), as candidate LPLCs. Notably, the emergence of specific LPLC markers, such as Sox9^5,16^ and EpCAM^25^, is proposed to result from injury-induced dedifferentiation of mature hepatocytes^26^. Considering the production of injury-responsive cells through dedifferentiation of mature cells via partial in vivo reprogramming in the intestine^13^, liver regeneration promoted by OSKM induction^12^ would likely result from the production of LPLCs through forced dedifferentiation in the absence of injury.

In this study, we demonstrated that forced dedifferentiation of the liver through partial in vivo reprogramming induced two distinct LPLC populations (LPLC1 and LPLC2) without triggering an immune response. This contrasts with LPLCs generated after injury, which are accompanied by a strong immune response. The LPLCs induced in the present study promoted liver regeneration following acute acetaminophen (N-acetyl-*p*-aminophenol: APAP)-induced liver injury (AILI). To develop a feasible strategy for translational applications, we employed OSKM delivered via a liver-specific lipid nanoparticle (LNP)-based nucleoside-modified mRNA delivery platform (OSKM mRNA-LNP). Intravenous administration of OSKM mRNA-LNP efficiently induced hepatocyte dedifferentiation, resulting in the production of LPLCs and promoting liver regeneration after sub-lethal APAP administration. Our findings highlight the potential of OSKM mRNA-LNP-based in vivo reprogramming for clinical applications, offering a promising approach for developing regenerative medicine for acute liver injuries.

## Results

### Hepatocyte dedifferentiation via partial reprogramming

Based on the formation of revSC-like cells (also known as ‘intestinal injury-responsive cells’)^3^ by transient OSKM induction^13^, we aimed to determine if in vivo partial reprogramming with short-course OSKM induction in a reprogrammable mouse model^27^ could produce LPLCs similar to the ‘hepatic injury-responsive cells’ in the absence of injury. A recent study reported hepatic and intestinal failure followed by premature death post-OSKM induction^28^; therefore, the hepatic condition was monitored closely during the OSKM induction process.

Previous studies have indicated that mice carrying a single doxycycline (Dox)-inducible OSKM cassette (4F^Tg/+^ and rtTA^Tg/+^) suffered premature death within three days when exposed to 1 mg/mL of Dox^28,29^. To avoid this complication, we employed a lower Dox dosage (0.15 mg/mL with 5% sucrose in drinking water) in homozygous mice (4F^Tg/Tg^ and rtTA^Tg/Tg^) for in vivo reprogramming as previously described^13^. This reprogramming condition caused a notable increase in OSKM expression within three days post-Dox treatment (Fig. 1a), accompanied by a marked expression of Oct4 in the liver (Fig. 1b, c).

**Figure 1.**
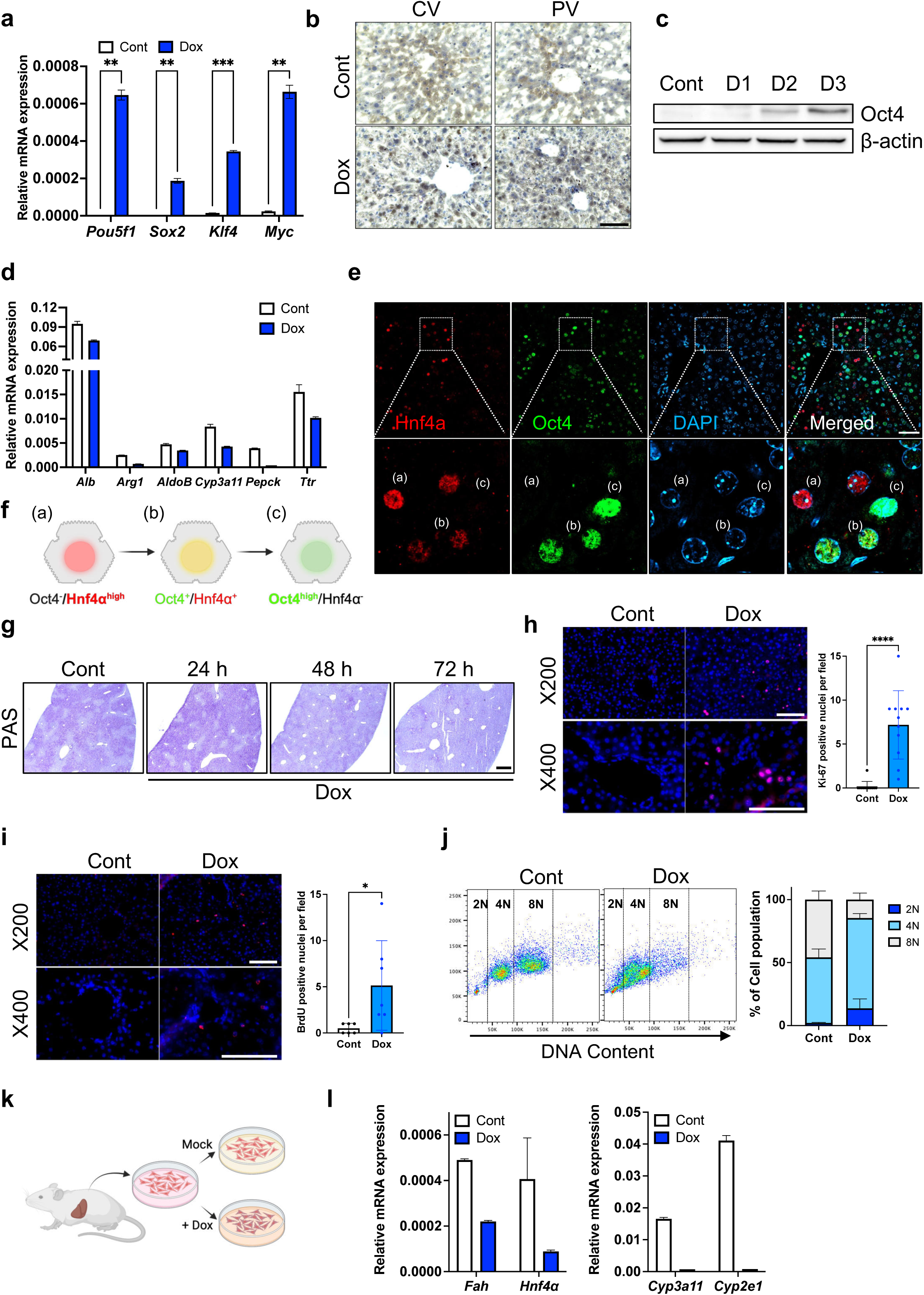
Hepatocyte dedifferentiation by partial reprogramming. (a) Relative mRNA expression of OSKM in isolated hepatocytes from OSKM inducible (iOSKM) mice at 3 days after Dox treatment. Data represent the mean with SD (n = 3) (b) Immunohistochemistry of Oct4 adjacent to central vein or portal vein. Samples were analyzed at 3 days after Dox treatment. Hematoxylin counterstaining were conducted. Scale bar = 100 μm (c) Immunoblot of Oct4 in whole liver lysates extracted at indicated days after Dox treatment. (d) Relative mRNA expression of hepatocyte-specific genes in isolated hepatocytes from iOSKM mice 3 days after doxycycline administration. Data represent the mean with SD (e) Immunofluorescence of Hnf4α (Red) and Oct4 (Green) in liver. DAPI (Blue) stains the nuclei. Samples were examined 48h after Dox treatment. (a) Hnf4α^high^/Oct4^-^ (b) Hnf4α^+^/Oct4^+^ (c) Hnf4α^-^/Oct4^high^ Scale bar = 100 μm (f) Schematic representation of hepatocyte dedifferentiation during partial reprogramming corresponding to Figure 1E. (g) PAS staining of liver which sampled at indicated days after Dox treatment. The purple area indicates cellular glycogen. Scale bar = 1 mm (h-i) Immunofluorescence of Ki-67 and BrdU (Red) in liver. DAPI (Blue) stains the nuclei (left). Samples were examined at 3 days after Dox treatment. Quantification of proliferating hepatocytes per field (n = 3, 2 ∼ 4 images per mice were analyzed) (right). Scale bar = 100 μm (j) Representative FACS plot of hepatocyte polyploidy (left) and the fraction of the 2n, 4n and 8n cells gated in the left panel (right). (n = 3) Data represent the mean with SD. (k) Schematic representation of in vitro partial reprogramming of isolated hepatocytes from iOSKM mice. (l) Relative mRNA expression of *Fah, Hnf4α, Cyp3a11, and Cyp2e1.* Isolated hepatocytes were cultured for 72 h under conditions with or without 1 μg/ml doxycycline. Statistical analysis was performed using a Student’s t-test: p < 0.05(*), p <0.01(**), p <0.001(***), p < 0.0001(****).

Importantly, this short-term induction protocol was well tolerated, showing no signs of weight loss (Supplementary Fig. 1a) or liver dysfunction, as evidenced by stable alanine aminotransferase (ALT) and aspartate aminotransferase (AST) during Dox administration (Supplementary Fig. 1b). Moreover, the levels of cleaved caspase 3 (cCasp3), which is associated with liver failure during in vivo reprogramming^28^, were minimal following the Dox treatment compared to control levels (Supplementary Fig. 1c). Notably, cCasp3 was absent throughout the liver after Dox treatment, in contrast to the notable increase in cCasp-positive areas near the central vein (CV) post-acetaminophen-induced liver injury (AILI) (Supplementary Fig. 1d). Hepatocytes near the CV, rather than those adjacent to the portal vein (PV), are exposed to a toxic metabolite of APAP (N-acetyl-p-benzo-quinone imine: NAPQI), which is synthesized due to the pericentral expression of the P450 enzyme^30^. Histological examinations conducted up to four days post-Dox treatment (0.15 mg/mL) revealed no structural changes or fibrosis (Supplementary Fig. 1e).

While no distinct deleterious effects were observed, temporary OSKM expression likely induced clear hepatic dedifferentiation. This was evidenced by the substantial decrease in hepatocyte-specific genes (Fig. 1d) and the reduced number of hepatocyte nuclear factor 4α (Hnf4α)-positive cells (Fig. 1e). Notably, immunofluorescence analysis revealed three distinct hepatocyte subpopulations (a) Oct4-/Hnf4α^+^ mature hepatocytes, (b) Oct4^+^/Hnf4α^low^ cells and (c) Oct4^high^/Hnf4α^-^ cells (Fig. 1e, f), suggesting a reprogramming-induced dedifferentiation. Additionally, the notable reduction in glycogen levels, evidenced by Periodic acid-Schiff staining (PAS) (Fig. 1g), indicated functional changes in the liver due to dedifferentiation. Similar to the increase in transient amplifying (TA) cells observed in the intestine post-OSKM induction^13^, we noted an increase in proliferating cells within the liver. This was demonstrated through Ki-67 staining (Fig. 1h) and BrdU incorporation in a pulse-chase experiment (Fig. 1i). Interestingly, while adult mouse hepatocytes are typically dominated by tetraploid (4N) and octoploid (8N) populations, hepatocytes from mice subjected to OSKM induction showed a marked reduction in the octoploid population and a corresponding increase in diploid (2N) cells (Fig. 1j). This shift in ploidy mirrors the profile observed in neonatal livers^31^ and in regenerating adult livers, suggesting that OSKM induction may promote a more proliferative, less terminally differentiated hepatic state. Further investigation involved isolating hepatocytes from adult mice and exposing them to Dox to assess the specific impact of OSKM on these cells (Fig. 1k). Consistently, we observed marked OSKM induction four days after Dox treatment (Supplementary Fig. 1f), coinciding with a decrease in markers associated with mature hepatocytes (Fig. 1l). Notably, the population of Oct4^high^/Hnf4αL cells—representing dedifferentiated hepatocytes—was markedly reduced beginning 48 hours after Dox withdrawal (Supplementary Fig. 1g), suggesting that in the absence of continued reprogramming stimulus, these cells rapidly redifferentiate into mature hepatocytes.

### Enhanced hepatic regeneration via partial reprogramming

Having successfully achieved hepatocyte dedifferentiation through partial reprogramming, we next focused on determining if this reprogramming could enhance liver repair after AILI, as demonstrated in a previous study^12^. To examine this, mice were challenged with a sublethal or lethal dose of APAP (300 mg/kg or 500 mg/kg, respectively), followed by Dox administration for three days to induce OSKM reprogramming. The animals were then monitored until Day 4 post-injury for assessment of regenerative outcomes (Fig. 2a).

**Figure 2.**
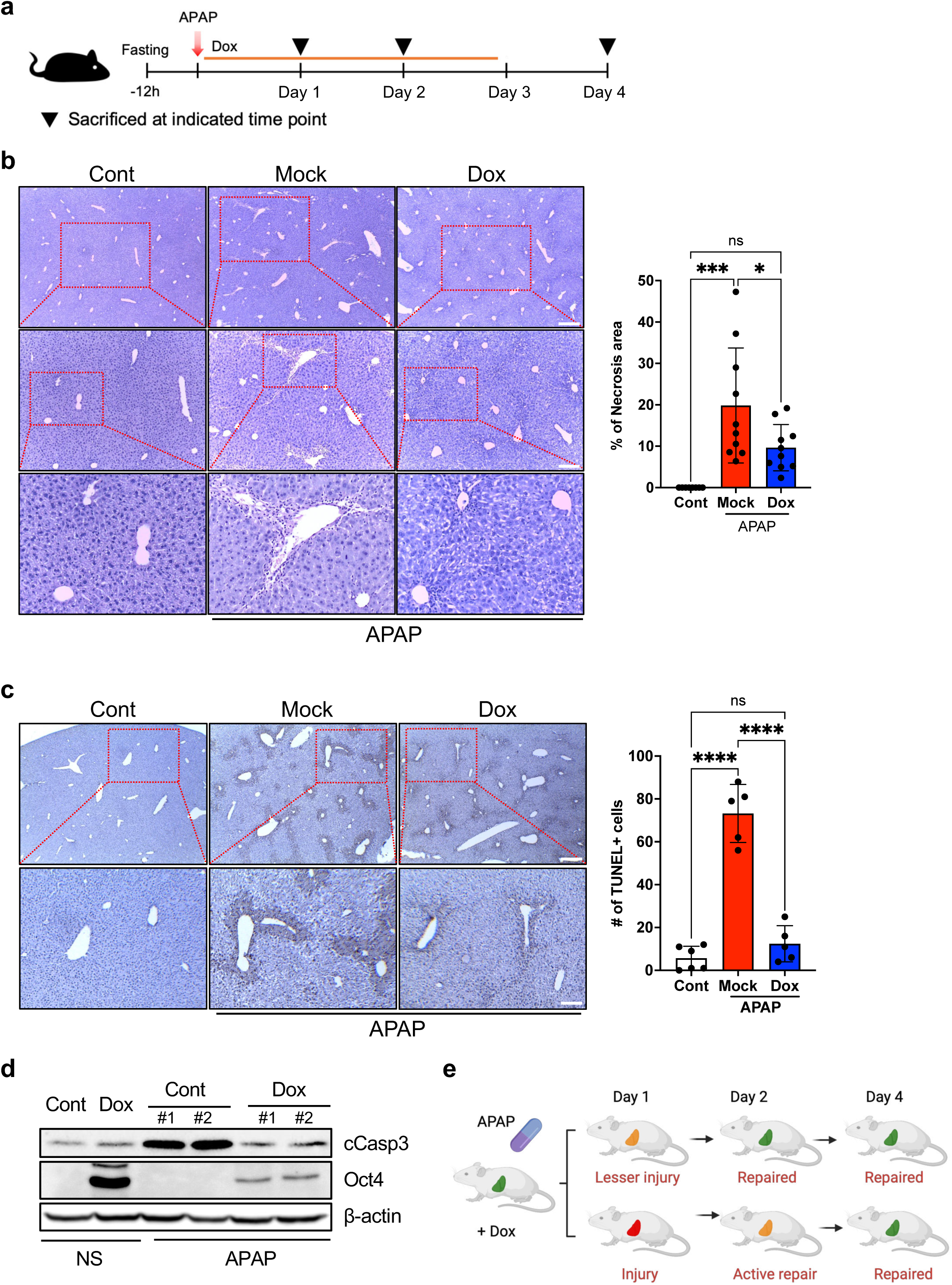
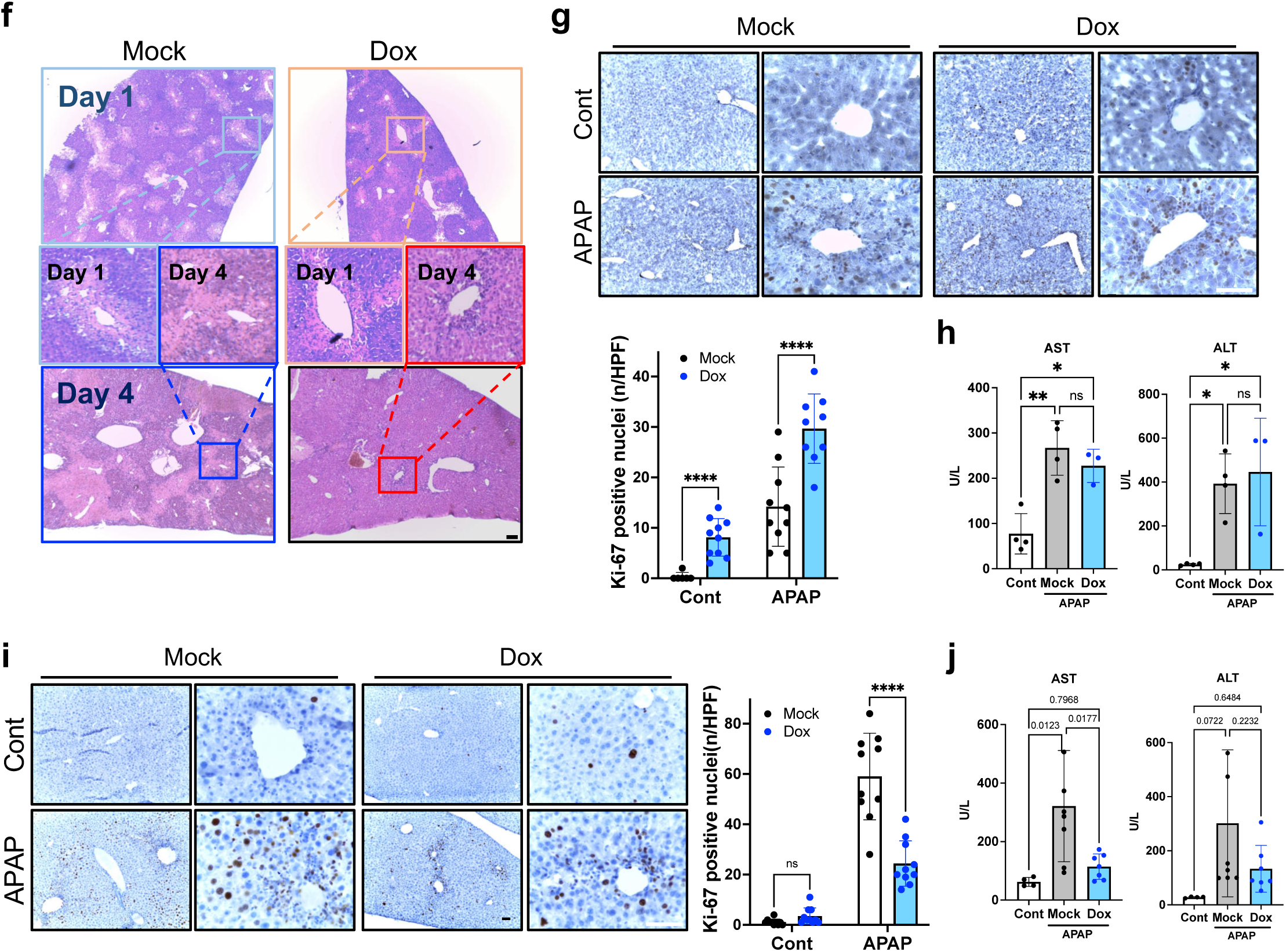
Enhanced hepatic regeneration by partial reprogramming. (a) Schematic illustration of partial reprogramming in an acetaminophen-induced liver injury (AILI) mouse model. iOSKM mice received intraperitoneal injection of APAP at a dose of 300 or 500 mg/kg after fasting for 12 hours. 0.15 mg/ml Dox treatment was followed immediately after APAP injection. (b) H&E staining of liver section sampled at 96 h post-AILI (500 mg/kg dose). Dotted line marks the necrotic area. (left) Percentage of necrosis area (right) (n=6, 1∼2 images per mice were analyzed). Scale bar = 250 μm (40x magnification), 100 μm (100x magnification) (c) TUNEL assay of liver section sampled at 96 h post-AILI (500 mg/kg dose). Images show the extent of DNA fragmntation, a hallmark of apoptosis. (left) Number of TUNEL positive cells per field (right) (n=6). Scale bar = 250 μm (40x magnification), 100 μm (100x magnification) (d) Immunoblot of active caspase-3 (cCasp3) and Oct4 in whole liver lysates extracted 48 h post-AILI (300 mg/kg dose). NS; Normal saline injected group, APAP; APAP injected group (e) Schematic illustration of putative mechanism for enhanced liver regeneration by partial reprogramming (f) H&E staining of liver section sampled at 24 h and 96 h post-AILI (500 mg/kg dose). Scale bar = 1 mm (g) Immunohistochemistry of Ki-67 adjacent to central vein. Samples were analyzed at 48 h post-AILI (300 mg/kg dose). Hematoxylin counterstaining were conducted. (upper) Quantification of Ki-67 positive nuclei per field (lower) (n = 3, 3 ∼ 4 images per mice were analyzed). Scale bar = 50 μm (h) Biochemical assays for serum ALT and AST levels (n = 3 ∼ 4) 48 h post-AILI (300 mg/kg dose). Data represent the mean with SD. Cont; normal mice, Mock; APAP (300 mg/kg I.P. injection) treated mice, Dox; APAP (300 mg/kg I.P. injection) treated + Doxycycline (0.15 mg/mg in drinking water) treated mice (i) Immunohistochemistry of Ki-67 adjacent to central vein. Samples were analyzed at 96 h post-AILI (500 mg/kg dose). Hematoxylin counterstaining were conducted. (left) Quantification of Ki-67 positive nuclei per field (right) (n = 3, 3 ∼ 4 images per mice were analyzed). Scale bar = 50 μm (j) Biochemical assays for serum ALT and AST levels (n = 4 ∼ 6) 96 h post-AILI (500 mg/kg dose). Data represent the mean with SD. Cont; normal mice, Mock; APAP (500 mg/kg I.P. injection) treated mice, Dox; APAP (500 mg/kg I.P. injection) treated + Doxycycline (0.15 mg/mg in drinking water) treated mice Statistical analysis was performed using two-way ANOVA (Fig. 2g, i) or one-way ANOVA (Figs. 2b, c, h, j): p < 0.05(*), p <0.01(**), p < 0.0001(****), ns, not significant.

At 96 hours post-AILI, we observed a marked reduction in both the necrotic area (Fig. 2b) and hepatocellular death, as evidenced by decreased TUNEL-positive cells (Fig. 2c) and reduced cleaved caspase-3 (cCasp3) levels (Fig. 2d) in the livers of mice subjected to OSKM induction. These findings were accompanied by the absence of hepatic fibrosis (Supplementary Fig. 2a) and reduced levels of 4-hydroxynonenal (4-HNE), a marker of lipid peroxidation and oxidative stress^32^ (Supplementary Fig. 2b). Considering the marked reduction in *Cyp2e1* and *Cyp3a11*—key enzymes involved in the bioactivation of APAP to the toxic metabolite NAPQI—following three days of OSKM induction (Fig. 1l), it became important to determine whether the observed reduction in hepatic damage was primarily due to decreased injury severity or enhanced regenerative activity (Fig. 2e). Histological analysis on Day 1 revealed comparable levels of necrosis between groups (Fig. 2f), and gross liver appearance, characterized by paleness and mottling indicative of necrosis and hemorrhage, also showed no apparent difference regardless of OSKM induction (Supplementary Fig. 2c). However, by Day 4 post-injury, OSKM-induced mice exhibited markedly improved liver morphology, suggesting enhanced tissue repair (Fig. 2f and Supplementary Fig. 2c). To distinguish enhanced regeneration from reduced initial injury, we closely monitored liver regeneration on Day 2 following moderate AILI. At this time point, OSKM-induced livers showed a significant increase in Ki-67-positive cells (Fig. 2g), despite comparable levels of liver damage as reflected by similar serum AST and ALT levels (Fig. 2h). In contrast, at Day 4 following severe AILI, proliferative activity was observed primarily in the mock group (Fig. 2i), coinciding with persistently elevated AST levels (Fig. 2j), indicative of unresolved injury. Despite the marked repression of *Cyp2e1* and *Cyp3a11* observed 24 hours after OSKM induction (Fig. 1l), the expression levels of these genes (Supplementary Fig. 2d) and the extent of initial liver injury (Supplementary Fig. 2e) remained comparable between groups within the first 6 hours after APAP administration—the critical window during which NAPQI production peaks^33^. In addition, high-resolution whole-liver images from mice subjected to APAP injury alone or APAP injury with OSKM induction clearly demonstrated a substantial increase in the Ki-67L population, accompanied by a loss of Hnf4α expression in the OSKM group, compared to control (Supplementary Data 1-3). These findings collectively support the conclusion that hepatic partial reprogramming significantly accelerates the liver repair process, consistent with previous reports^12^.

### Generation of two distinct progenitor-like populations via OSKM induction

We previously demonstrated that partial reprogramming produces ‘injury-responsive cells’ by recapitulating dedifferentiation patterns observed following intestinal injury^13^. Given the clear hepatic dedifferentiation elicited by partial reprogramming (Fig. 1), we hypothesized that this process may be sufficient to give rise to ‘injury-responsive cells’, specifically LPLCsL, independent of exogenous injury. To investigate this hypothesis, we performed single-nucleus RNA sequencing (snRNA-seq) on livers subjected to OSKM-mediated reprogramming and compared these profiles with those obtained from livers following APAP-induced injury. Leveraging our established computational pipeline for quality control and batch correction, we constructed a high-resolution atlas of the partially reprogrammed hepatic environment, encompassing 22,641 quality-filtered nuclei, with an average of 1,642 detected genes and 3,087 unique molecular identifiers (UMIs) per cell (Fig. 3a, left). Unsupervised clustering and canonical marker gene expression analysis identified eight major hepatic cell types (Supplementary Fig. 3a, left), including hepatocytes, Kupffer cells (KCs), monocytes/monocyte-derived macrophages (Mono/MoMq), hepatic stellate cells (HSCs), endothelial cells (ECs), mesothelial cells, T cells, and B cells. (Supplementary Fig. 3a, left). A prominent difference between the APAP and OSKM conditions was the substantial increase in immune cell populations, including KC and Mono/MoMq, observed predominantly in the APAP-injured liver (4.5- and 16-fold increases compared to the control, respectively). In contrast, these populations were relatively limited in the OSKM-induced liver (Fig. 3a, right). These findings suggest that liver reprogramming triggers a marginal immune response compared to AILI. To further investigate hepatocyte dynamics, we subdivided hepatocyte populations based on established marker gene expression (Supplementary Fig. 3a, right) allowing us to assess the effects of both APAP injury and OSKM-mediated reprogramming (Fig. 3b). Notably, pericentral hepatocytes (Hep_CV) showed a pronounced reduction in relative abundance following APAP-induced injury. Intriguingly, reductions in both Hep_CV and periportal hepatocytes (Hep_PV) were also observed after OSKM induction, despite the absence of overt cell death (Fig. 3c) as seen in the Supplementary Fig. 1d.

**Figure 3.**
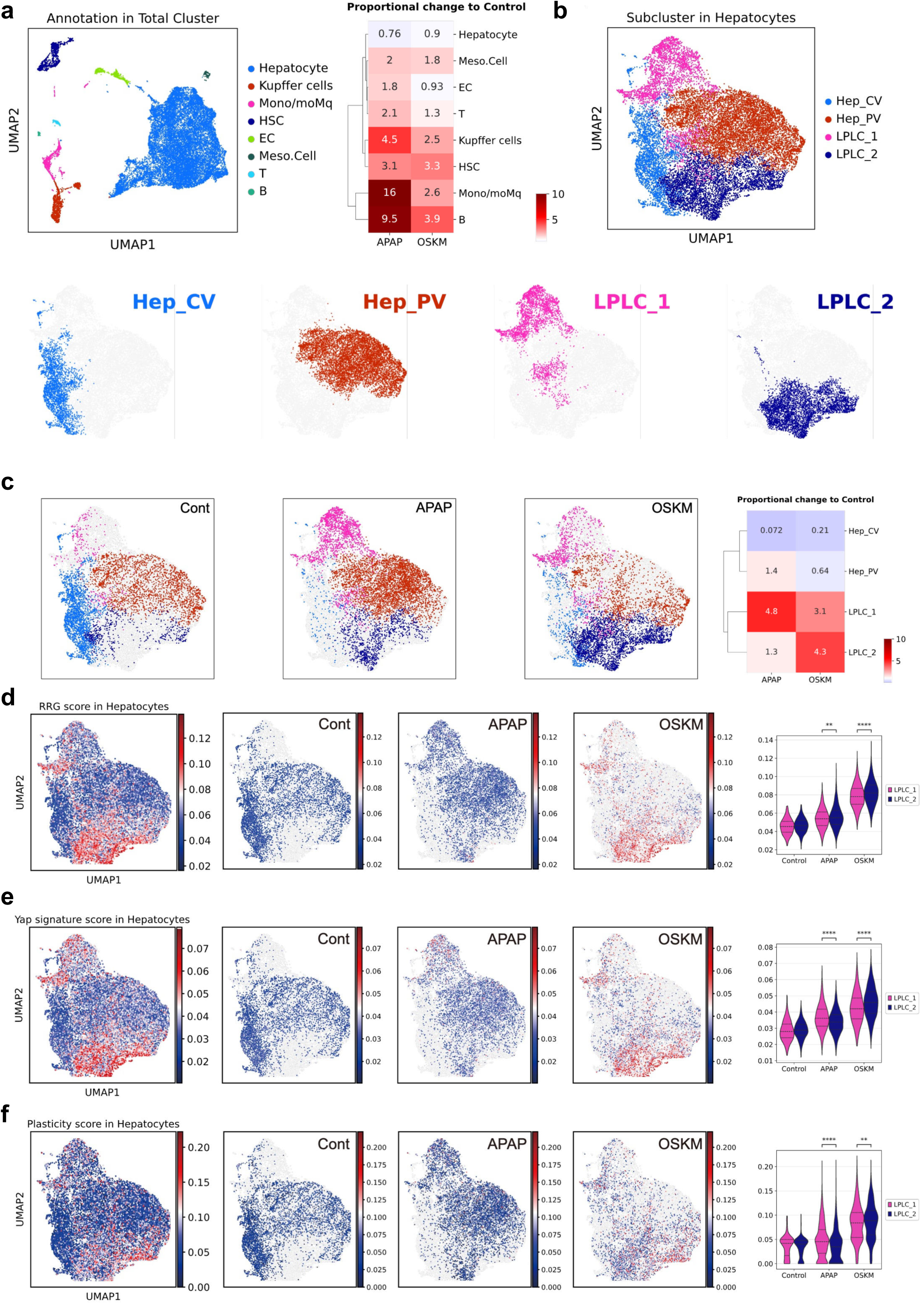
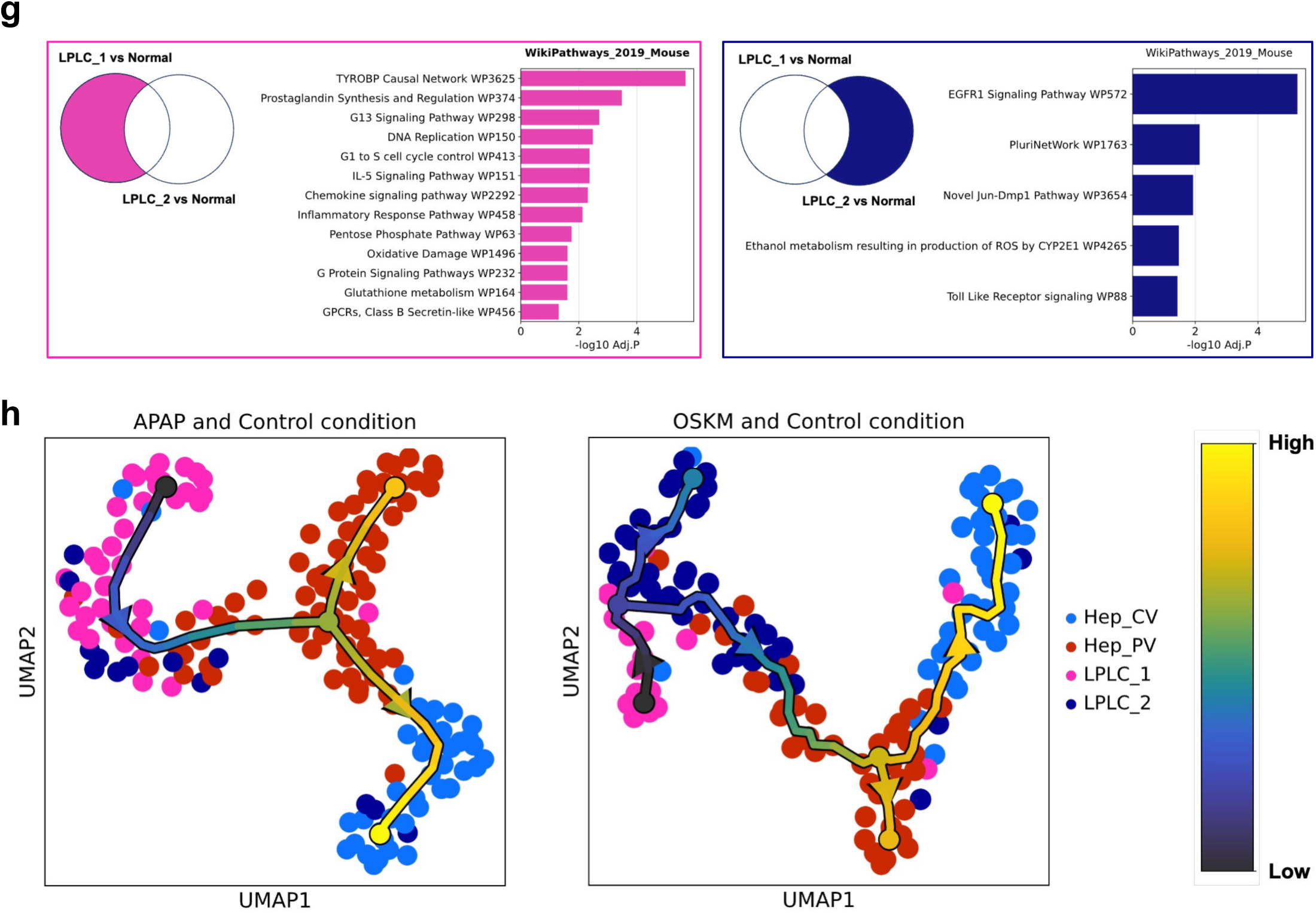
Identification of distinct two LPLC populations induced by partial reprogramming. (a) UMAP visualization of liver cell clusters from single nucleus RNA sequencing (snRNA-seq) results (left) and relative fold change of proportion compared to Control condition (right). Each value in clustermap represents relative fold change of cell type proportion to Control condition. (b) UMAP visualization of hepatocytes (upper right). Each UMAP indicates distribution of cell types in hepatocytes (lower). (c) Distribution of cell types in each condition (left) and relative fold change to Control condition in Hepatocyte subpopulation (right) (d-f) Reprogramming/progenitor-related genes (RRGs) score, YAP signature score, and Plasticity signature score values were represented on Hepatocyte UMAP (left) and violin plot (right). The scores were calculated with ‘AUCell’. Gene sets used for scoring were from previous studies or Wikipathway (2019) ‘PluriNetWork’. (g) Upregulated terms specific to LPLC_1 or LPLC_2 were presented. (h) Trajectory analysis in hepatocytes with selected metacells. Each analysis only contains one of the case conditions (APAP or OSKM) and Control. The color gradient of colorbar (right) indicates pseudotime value.

Importantly, we identified two distinct subpopulations of liver progenitor-like cells (LPLCs), designated LPLC_1 and LPLC_2, under both conditions. In APAP-injured livers, LPLC_1 was markedly expanded (4.8-fold increase versus control), whereas both LPLC_1 and LPLC_2 were elevated in OSKM-induced livers (3.1- and 4.3-fold increases, respectively; Fig. 3c). These results indicate that partial reprogramming alone is sufficient to induce the emergence of LPLCs, highlighting their potential role in liver regeneration. We hypothesized that these injury- or reprogramming-induced LPLC subpopulations would exhibit heightened cellular plasticity. To evaluate this, we calculated reprogramming/progenitor gene (RRG) scores, which reflect the transcriptional features of injury-responsive LPLCs^5^. While LPLCs generated post-APAP injury showed increased RRG scores, both LPLC_1 and LPLC_2 demonstrated even higher scores following OSKM induction (Fig. 3d). This enrichment was further supported by elevated YAP signature (Fig. 3e) and plasticity scores (Fig. 3f) in both conditions, consistent with the known roles of these signatures in cellular dedifferentiation and regenerative capacity (Supplementary Data 4). Gene set enrichment analysis (GSEA) revealed functional distinctions between LPLC subtypes. While both LPLC_1 and LPLC_2 were enriched for genes involved in active cellular processes (Supplementary Fig. 3b), LPLC_1 was specifically associated with proliferative pathways, including ‘DNA replication’ and ‘cell cycle control’ (Fig. 3g, left). This suggests a key role for LPLC_1 in promoting hepatocyte cell cycle progression following injury or reprogramming. In contrast, LPLC_2 was enriched in the ‘PluriNetWork’ pathway (Fig. 3g, right), indicative of a dedifferentiated, highly plastic cellular state, likely induced by OSKM reprogramming (Fig. 3f). Among the well-established LPLC markers^5^, *Spp1* and *Cd44*, both implicated in hepatic regeneration^34^, were more highly expressed in LPLC_1 than in LPLC_2 (Supplementary Fig. 3c). To examine lineage potential, we performed SEACell-based metacell analysis and trajectory inference, comparing control with either APAP or OSKM conditions. This revealed that LPLC_1 occupies a central position along the predicted differentiation trajectories and likely serves as a progenitor for regenerating both Hep_CV and Hep_PV populations under both injury and reprogramming conditions (Fig. 3h). Collectively, these findings support a model in which LPLC_1 is a key cellular intermediary in liver regeneration, activated by both APAP-induced injury and OSKM-mediated partial reprogramming.

### Zonation-specific LPLC formation following AILI or partial reprogramming

For further characterization of LPLCs, we aimed to determine the specific timeframe in which hepatocyte dedifferentiation is initiated in response to injury and to identify their unique transcriptomic signature. Using publicly available RNA sequencing (RNAseq) data with whole liver samples collected at multiple time points post-AILI^35^ (accessible at https://zenodo.org/records/6035873), we monitored the time-dependent transcriptome profile. Principal component analysis (PCA) revealed significant transcriptomic alterations by 48 h post-AILI, with a gradual reversion to normal liver transcriptomic patterns by 96 h post-injury (Fig. 4a). We hypothesized that stress (or damage) genes associated with cell death and inflammatory responses would be upregulated immediately after injury. This would be succeeded by genes involved in ’repair, dedifferentiation, or regeneration’, with those related to ‘hepatic function’ recovering subsequently (Fig. 4b). As anticipated, gene signatures indicative of stress and damage responses, including ‘p53 pathway’, ‘apoptosis’, and ‘inflammatory response’ exhibited high normalized enrichment scores (NES) at 6 h post-AILI, followed by a decline over time (Fig. 4c, red). In contrast, the NES for cell cycle genes, such as ‘E2F targets’ and ‘G2/M checkpoint’, peaked at 48 h (Fig. 4c, green). Notably, the NES for dedifferentiation genes, such as ‘reprogramming-related genes’ and ‘negative regulation of epithelial differentiation’, was evident at 24 h and then gradually decreased (Fig. 4c, blue).

**Figure 4.**
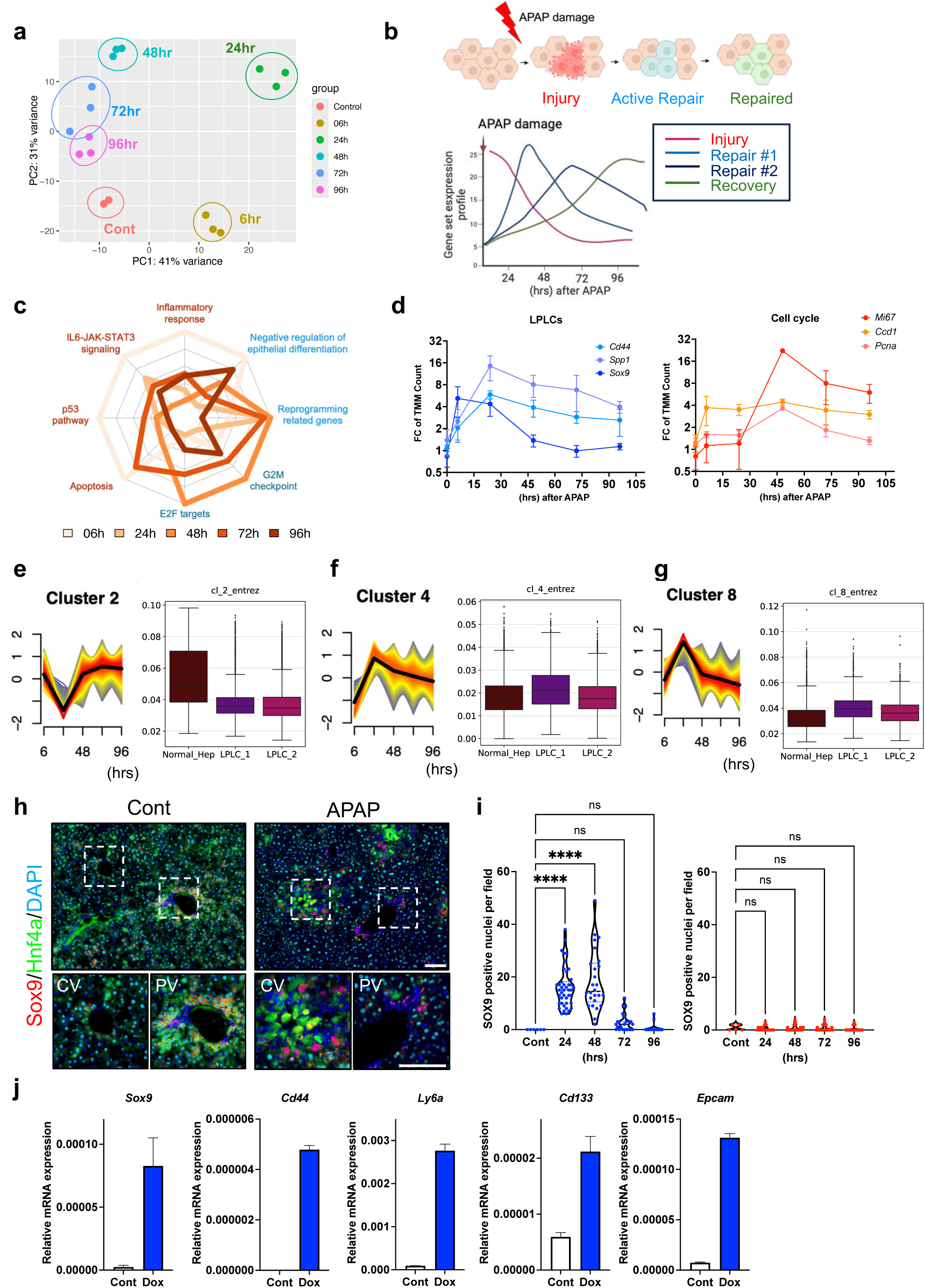
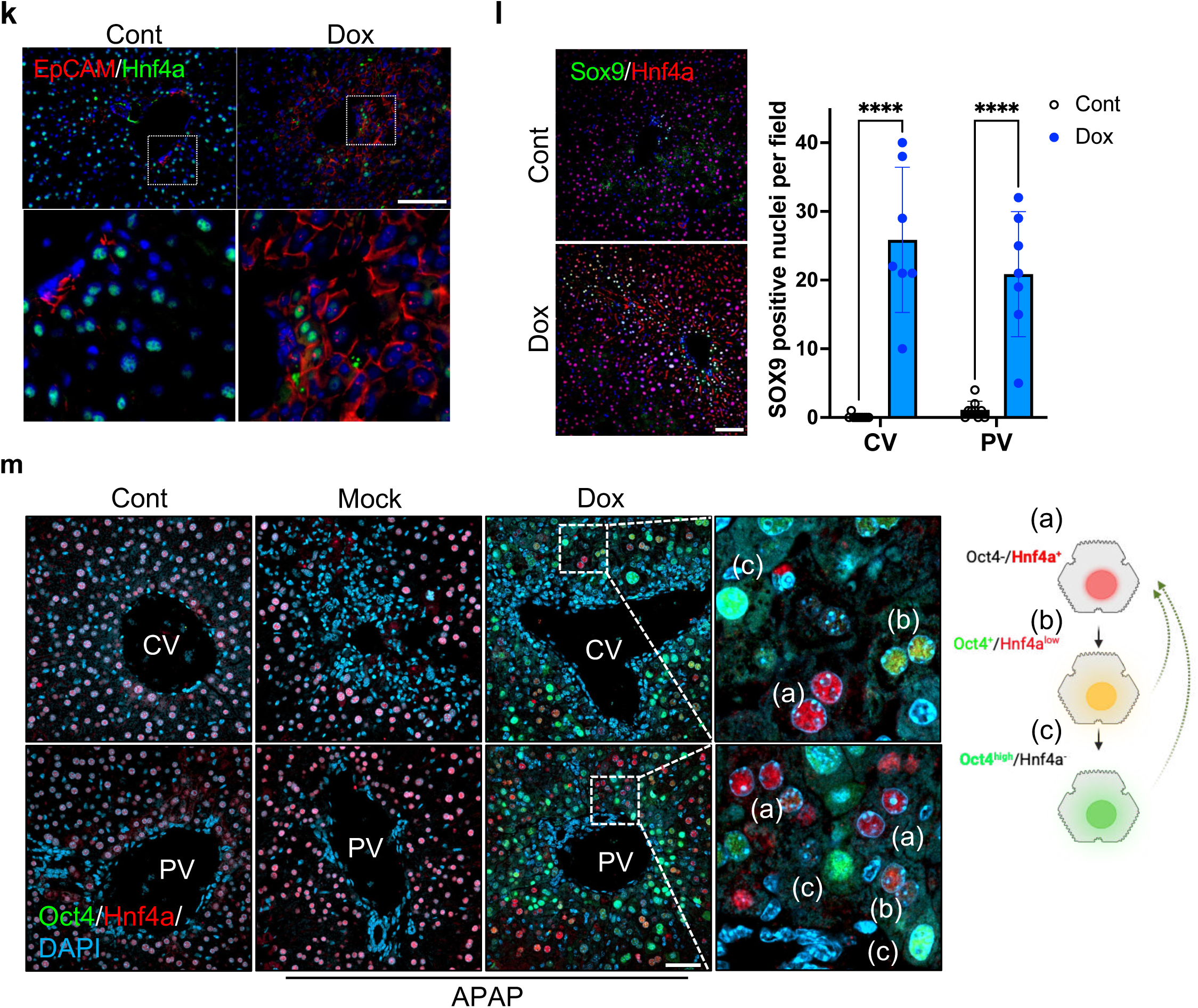
Partial reprogramming recapitulates hepatic regeneration process after AILI. (a) Principal component analysis (PCA) of liver post-AILI. Control (Orange), 6 h (Yellow), 24 h (Green), 48 h (Cyan), 72 h (Blue), 96 h (Purple) were analyzed. (b) Schematic illustration of time-course transcriptomic alteration through the regeneration process. (c) Radar plot for normalized enrichment score (NES) of time-course GSEA with indicated hallmark genes. (d) Time-course fluctuations of trimmed mean of M values (TMM count) for visualizing LPLC / Cell cycle marker expression patterns. (e-g) Module scores of cluster-specific genes (left) using hepatocyte subcluster (Fig. 3) were represented on box plots (right). (h) Immunofluorescence of Sox9 (Red) and Hnf4α (Green) in liver sections sampled 0 h / 48 h post-AILI (300 mg/kg dose). DAPI (Blue) stains the nuclei. Insets indicate pericentral area and periportal area. Scale bar = 100 μm (i) Quantification of Sox9 positive nuclei per field adjacent to CV (left, blue) and PV (right, red). (n=3, 3∼14 images per mice were analyzed) (j) Relative mRNA expression of LPLC markers listed in Supplementary Fig. 4d in isolated hepatocytes from iOSKM mice at 3 days after Dox treatment. (k) Immunofluorescence of EpCAM (Red) and Hnf4α (Green) in liver sections from control and Dox treated iOSKM mouse. DAPI (Blue) stains the nuclei. Scale bar = 100 μm. (l) Immunofluorescence of Sox9 (Green) and Hnf4α (Red) in liver sections from control and Dox treated iOSKM mouse. DAPI stains the nuclei (left). The number of Sox9 positive nuclei per field adjacent to CV and PV (n=3, 2 ∼ 4 images per mice were analyzed) (right). (m) Immunofluorescence of Oct4 (Green) and Hnf4α (Red) adjacent to CV and PV. DAPI (Blue) stains the nuclei. (right) Schematic representation of putative mechanism of redifferentiation after partial reprogramming (left). Cont; normal mice, Mock; APAP (300 mg/kg I.P. injection) treated mice, Dox; APAP (300 mg/kg I.P. injection) treated + Doxycycline (0.15 mg/mg in drinking water) treated mice. Scale bar = 50 μm. Statistical analysis was performed using one-way ANOVA: p < 0.0001(****), ns, not significant.

Temporal enrichment of gene sets related to ‘dedifferentiation’ and ‘cell cycle’ was highlighted by the time-dependent expression profiles of key representative genes post-AILI (Fig. 4d). Unbiased soft clustering technique grouped the genes based on their expression profiles over time, revealing eight specific time-course patterns (Supplementary Fig. 4a). Cluster 2, representing the recovery gene profile (Fig. 4b), was specifically distinct in normal hepatocytes (Fig. 4e). In contrast, clusters 4 and 8, analogous to the repair profiles (Fig. 4b), were enriched in LPLC_1 (Fig. 4f, g). Additionally, we observed a significant increase in Sox9-positive LPLCs (Sox9+ LPLCs) at both 24- and 48-h post-AILI (Supplementary Fig. 4c). Notably, these Sox9+ LPLCs were primarily localized in the pericentral area—the primary site for AILI^36^—but not in the periportal area (Fig. 4h, i). The formation of LPLC_1, similar to that observed in AILI, was validated under OSKM induction (Fig. 3). This was evidenced by the upregulation of typical LPLC marker genes, including *Sox9, Cd44, Ly6a, Cd133*, and *Epcam (*Fig. 4j and Supplementary Fig. 4d), as well as the protein expression of EpCAM, a marker for hepatic stem/progenitor cells^37^ or newly derived hepatocytes^38^, alongside the repression of the Hnf4α protein (Fig. 4k). In sharp contrast to the zonation-specific LPLC induction observed post-AILI, highlighted by co-staining with glutamine synthetase (GS), which is exclusively expressed in CV hepatocytes (Supplementary Fig. 4e), OSKM induction resulted in a marked increase in the Sox9L population (Fig. 4l) along with dedifferentiated hepatocytes identified as (b) Oct4^+^/Hnf4α^low^ and (c) Oct4^high^/Hnf4 ^-^ (Fig. 4m) in both CV and PV regions. Given the Oct4-dependent repression of Hnf4α shown in Fig. 1E, and the time-dependent re-expression of Hnf4α following Dox withdrawal (Supplementary Fig. 1g), the presence of (a) Oct4L/Hnf4αL, (b) Oct4^+^/Hnf4α^low^ and (c) Oct4^high^/Hnf4α^-^ cells across both Hep_CV and Hep_PV populations in the single-cell transcriptomic data (Fig. 3c) supports the occurrence of widespread hepatocyte dedifferentiation across hepatic zones following OSKM induction.

### Immune response-free dedifferentiation via partial reprogramming

Injury-induced immune responses typically provoke cellular dedifferentiation, producing ‘injury-responsive cells’ specific to each organ, orchestrated by cytokines and signaling molecules such as IL-6^5^, TGF-b^6^, and PGE2^7^. Notably, autonomous production of PGE2 facilitates intestinal repair after partial reprogramming without involving the immune response^13^. We investigated whether this immune-independent reprogramming also occurs in the liver, mirroring the intestinal process (Fig. 5a). Interestingly, Stat3 phosphorylation—a key downstream signal of IL-6—along with Sox9 induction and Hnf4a repression, was evident in primary hepatocytes post-OSKM induction; however, these changes were not observed in the absence of the OSKM cassette (Fig. 5b), indicating that OSKM-mediated reprogramming alone is sufficient to activate this dedifferentiation program. Time-course analysis revealed that both APAP-induced liver injury (AILI) and OSKM induction elicited similar temporal patterns of Stat3 phosphorylation and Sox9 expression, suggesting that OSKM-induced LPLC formation recapitulates the molecular trajectory of injury-driven dedifferentiation (Fig. 5c).

**Figure 5.**
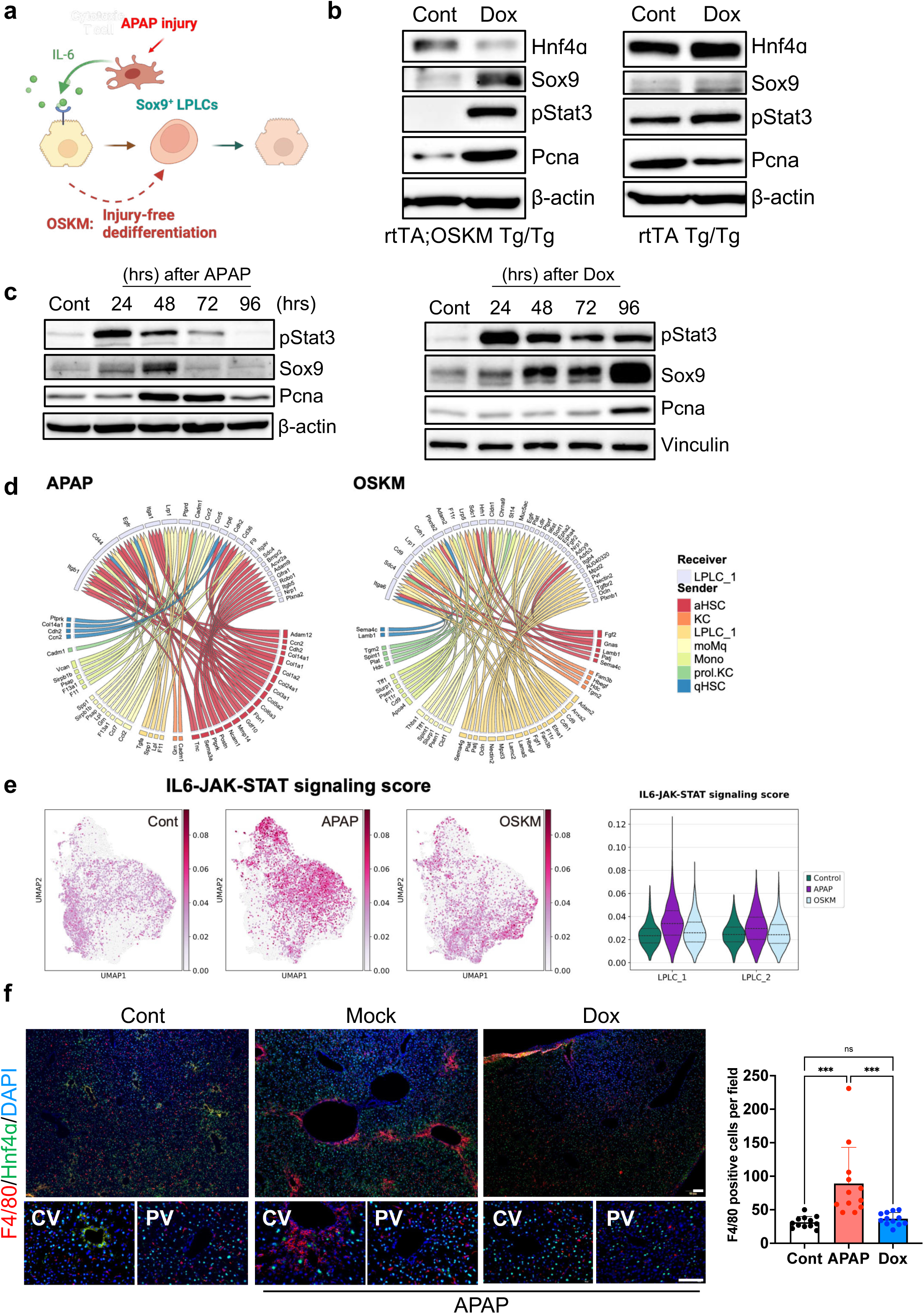
Inflammation-free hepatocyte dedifferentiation by partial reprogramming. (a) Schematic illustration of putative mechanism for hepatocyte dedifferentiation by partial reprogramming. (b) Immunoblot of Hnf4α, Sox9, pStat3 and PCNA in isolated hepatocytes from iOSKM mice or ROSA26-rtTA^Tg/Tg^ mice. All hepatocytes were isolated 3 days after Dox treatment. (c) Immunoblot of Sox9, pStat3 and PCNA in whole liver lysates sampled at indicated timepoint post-AILI (left). Immunoblot of Sox9, pStat3 and PCNA of whole liver lysates sampled at indicated timepoint after OSKM induction (right). (d) Results of cell-cell communication analysis using NicheNet. Lower semicircle indicates significantly upregulated ligands of each cell type. Upper semicircle indicates receptors of LPLC for each ligand. (e) IL6-JAK-Stat3 signaling score presented on UMAP of hepatocyte (left) and on violin plot of LPLC_1 and LPLC_2 (right). (f) Immunofluorescence of F4/80 (Red) and Hnf4α (Green) in liver sections. DAPI stains the nuclei (Blue). All samples were collected 48h post-injection. Cont; normal mice, Mock; APAP (300 mg/kg I.P. injection) treated mice, Dox; APAP (300 mg/kg I.P. injection) treated + Doxycycline (0.15 mg/mg in drinking water) treated mice. (left) Quantification of F4/80 positive cells per field (right). Scale bar = 100 μm Statistical analysis was performed using one-way ANOVA: p < 0.001(***), ns, not significant.

To explore the intercellular signaling landscape underpinning this process, we conducted NicheNet-based analysis of cell-to-cell communication under both AILI and OSKM^39^. Based on prior trajectory analysis (Fig. 3h), LPLC_1 was designated as the signal-receiving population. Potential ligand–receptor interactions were prioritized using a threshold interaction score >0.8 (Supplementary Fig. 5a). Under AILI conditions, a range of immune cell types emerged as candidate signal senders; however, activated hepatic stellate cells (aHSCs) exhibited the highest number of prioritized interactions (Fig. 5d, left). In contrast, under OSKM-induced conditions, most signals originated from LPLC_1 itself or HSCs (Fig. 5d, right). These findings suggest that liver regeneration following AILI is initiated by immune-derived signals, whereas OSKM-induced regeneration proceed largely independent of immune activation. Furthermore, gene set enrichment analysis demonstrated significant upregulation of the IL6–JAK–STAT signaling pathway in LPLC_1 cells post-AILI, with notably higher pathway activity scores in the APAP-injured condition (Supplementary Data 4). This result confirmed a pronounced enrichment of IL6–JAK–STAT signaling specifically in LPLC_1 following AILI, reinforcing the role of immune-derived cytokine signaling in injury-mediated, but not reprogramming-mediated, liver regeneration.

To validate these predictions, we examined the spatial distribution of LPLCs and immune cells. After AILI, LPLCs and F4/80-positive macrophages were predominantly localized near the injured area (Fig. 4l and 5f). Conversely, the level of F4/80-positive cells three days after OSKM induction was similar to that observed in the control group (Fig. 5f). These findings suggest that hepatic reprogramming through a transient OSKM challenge successfully generates LPLCs while bypassing the immune response typically observed in injury-induced scenarios. This process appears to contribute effectively to liver regeneration following AILI.

### Hepatocyte dedifferentiation via liver-targeted delivery of OSKM mRNA

To translate the previously demonstrated effects of OSKM induction to therapeutically relevant clinical settings, we utilized an LNP-based nucleoside-modified mRNA (LNP-mRNA) delivery system. Previous studies^9^ have employed recombinant adeno-associated virus (AAV) vectors for the ectopic expression of the OSK. However, prolonged OSKM induction poses significant risks, including the potential for intestine or liver failure^28^ and premature death^29^. Thus, continuous expression via AAV delivery over several weeks^40^ would lead to complications. In contrast, LNP-mRNA allows for highly reproducible, transient gene expression with a single administration^41^. Additionally, the duration of gene expression can be controlled through multiple administrations, making it safer for translational applications of OSKM compared to viral delivery. To specifically target the liver, we used a galactosylated formulation of C12-SPM ionizable lipid-based LNP (C12-SPM-GAL), as described in our previous study^42^. This formulation contained galactosylated ceramide as the fifth lipid component to target galactose receptors on hepatocytes^43^. We optimized the previous LNP-siRNA formulation into an LNP-mRNA formulation by replacing the helper lipid with 1,2-dioleoyl-sn-glycero-3-phosphoethanolamine (DOPE) and adjusting the ratios of the lipid components (Fig. 6a). When C12-SPM-GAL-based LNPs (GAL) loaded with firefly luciferase (Luc) mRNA were injected intravenously into mice at an mRNA dose of 0.3 mg/kg, there was no significant difference in ALT and AST levels compared to the phosphate-buffered saline (PBS)-injected group, indicating that the LNPs induced minimal hepatotoxicity (Fig. 6b) and negligible cytotoxicity (Supplementary Fig. 6a). Compared to LNPs formulated with the FDA-approved ionizable lipid DLin-MC3-DMA (MC3)^41^, C12-SPM-GAL LNPs had minimal impact on serum ALT and AST levels (Fig. 6b) and demonstrated superior mRNA delivery efficacy in the liver, with maximal expression at 6 h post-injection and a gradual decrease over 24 h (Fig. 6c). Ex vivo bioluminescence of major organs at 6 hours post-injection revealed that C12-SPM-GAL LNP achieved significantly higher firefly luciferase expression in the liver compared to conventional LNPs (MC3 and SM-102), with approximately 98.9% of total firefly luciferase expression localized to the liver (Supplementary Fig. 6b, c). While serum AST, ALT, and IL-6 levels transiently increased in GAL group at 6 hours post-LNP administration, these changes were comparable to those observed with MC3 and SM-102 LNPs (Supplementary Fig. 6d) and returned to levels similar to PBS-injected group within 24 hours post-LNP administration (Fig. 6b). We then synthesized nucleoside-modified OSKM mRNAs and confirmed their correct synthesis (Supplementary Fig. 6e). The LNPs encapsulating the four factor mRNAs showed homogeneous characteristics, with over 85% encapsulation efficiency and particle sizes less than 100 nm (Supplementary Fig. 6f). Initial validation of protein expression using these LNPs was performed in mouse embryonic fibroblasts (MEF) (Fig. 6d and Supplementary Fig. 6g). Subsequently, OSKM mRNA-LNPs were introduced into mice via retro-orbital injection at an mRNA dose of 0.3 mg/kg, and the effects were monitored at 30- and 54-h post-administration (Fig. 6e). Since mRNA expression peaked as early as 6 h post-injection, mRNA levels in liver tissues significantly decreased by 54 h post-injection (Supplementary Fig. 6h). The reprogramming effect of OSKM mRNA-LNPs was verified by the induction of *Sox9* and *Ki-67* mRNAs (Fig. 6f) and their corresponding protein levels (Fig. 6g) at 30 h post-injection. The formation of Sox9+ LPLCs was accompanied by active cell cycling, as indicated by Histone H3 phosphorylation, a mitotic marker, and an increase in PCNA protein levels (Fig. 6h). Additionally, histological analysis revealed that OSKM mRNA-LNP treatment induced no detrimental effects in liver tissues at both indicated doses (Supplementary Fig. 6i). These findings demonstrate that the LNP-mRNA system enables efficient in vivo reprogramming through transient, liver-specific expression of OSKM.

**Figure 6.**
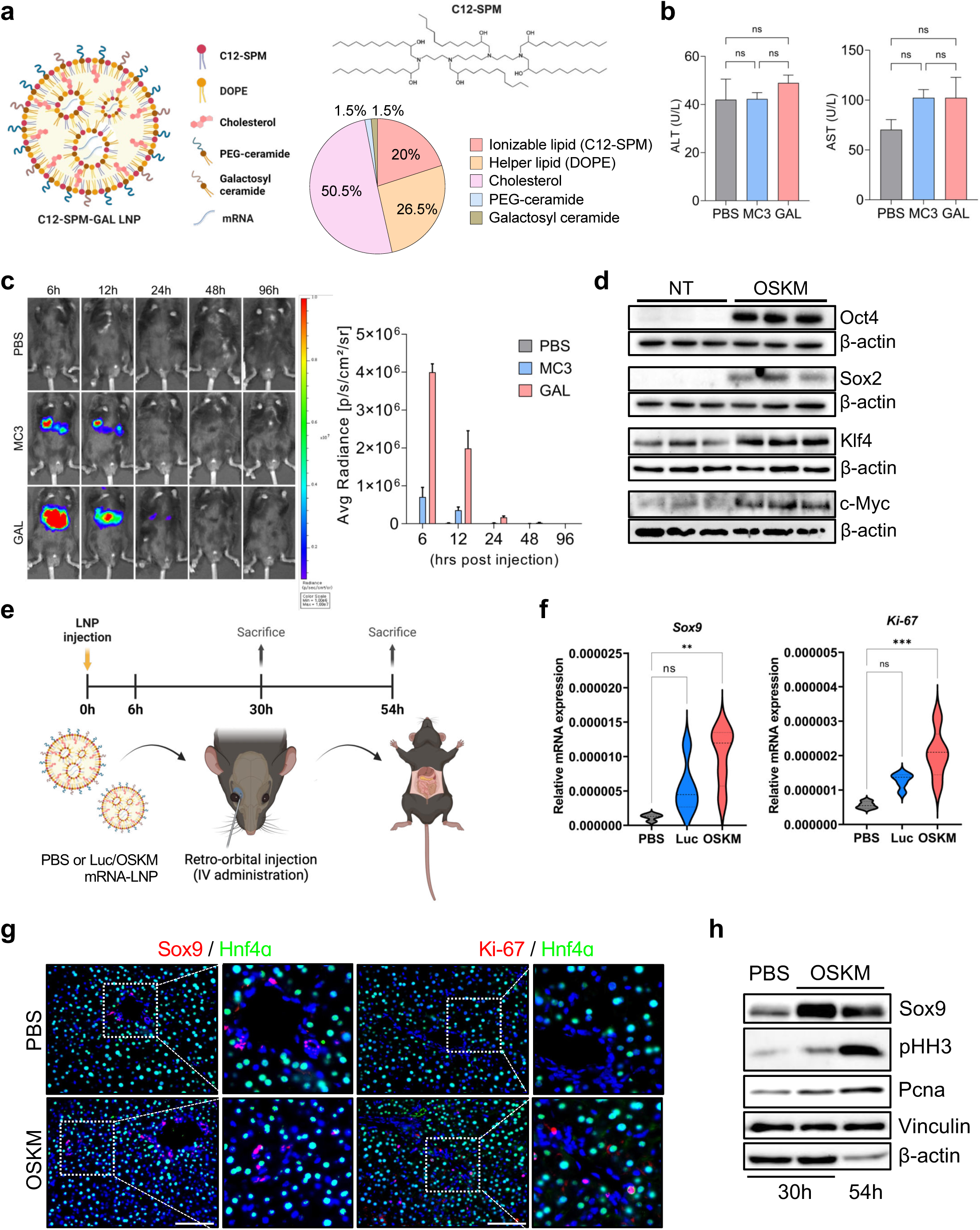
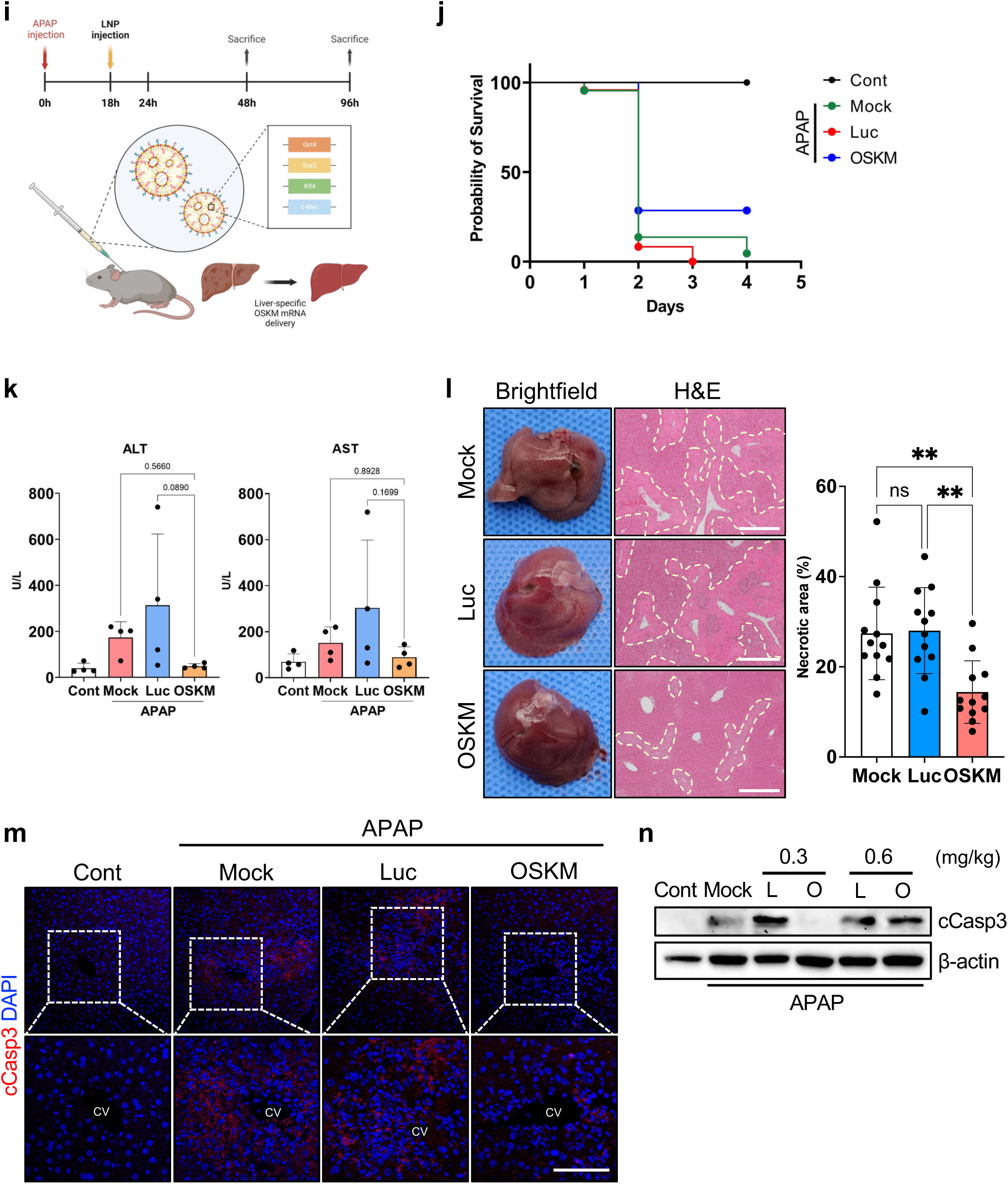
Hepatocyte dedifferentiation by liver-targeted delivery of nucleoside-modified OSKM-mRNA. (a) Illustration of C12-SPM-GAL LNP-mRNA formulation. Chemical structure of the ionizable lipid C12-SPM and the lipid components with their respective molar ratios in the optimized C12-SPM-GAL LNP-mRNA formulation. (b) Biochemical assays for serum ALT and AST levels (n = 3) 24 h after intravenous (IV) administration of LNPs loaded with firefly luciferase mRNA (0.3 mg/kg) in C57BL/6 mice. PBS; PBS-injected mice, MC3; DLin-MC3-DMA LNP-injected mice, GAL; C12-SPM-GAL LNP-injected mice. (c) Time-course analysis of in vivo bioluminescence following IV administration of LNPs loaded with firefly luciferase mRNA in C57BL/6 mice (0.3 mg/kg) (n = 3 ∼ 4). (d) Detection of OSKM proteins 24 h after transfection of C12-SPM-GAL LNPs loaded with OSKM mRNA into MEFs (1 μg total OSKM mRNA for 100,000 cells, 0.25 μg for each mRNA). NT; no treatment, OSKM; OSKM mRNA-LNP treatment. (e) Experimental outline for assessing in vivo partial reprogramming by OSKM mRNA-LNP to induce hepatocyte dedifferentiation in normal wild-type mice. The PBS-injected group (PBS group) was used as a negative control, and both Luc mRNA-LNP (Luc group) and OSKM mRNA-LNP (OSKM group) were IV administered at a dose of 0.3 mg/kg. (f) Quantitative PCR analysis of liver progenitor-like cell (LPLC)-related genes in the liver 30 h post-injection (n = 4 ∼ 5). (g) Immunofluorescence staining of Sox9 and Ki-67 co-stained with Hnf4α in the liver 30 h post-injection. DAPI stains the nuclei (scale bars = 100 μm). (h) Western blot analysis for LPLC marker Sox9, cell cycle marker phosphorylated histone H3 (pHH3), and proliferating cell nuclear antigen (PCNA) in the liver 30 and 54 h post-injection. (i) Schematic illustration of C12-SPM-GAL-mediated OSKM mRNA treatment in an acetaminophen (APAP)-induced acute liver injury mouse model. C57BL/6 mice received intraperitoneal injection of APAP at a dose of 400 mg/kg after fasting for 12 h. Mice were treated with PBS (Mock group), Luc mRNA-LNP (Luc group), or OSKM mRNA-LNP (OSKM group) 18 h post-APAP administration. (j) Kaplan-Meier survival plot of mice (n = 14) in each treatment group over four days following APAP administration. Cont; normal mice, Mock; APAP injury + PBS-injected mice, Luc; APAP injury + Luc mRNA-LNP-injected mice, OSKM; APAP injury + OSKM mRNA-LNP-injected mice. (k) Blood chemistry analysis of serum AST and ALT levels 96 hours post-APAP administration (78 hours post-treatment). (l) Representative images of the whole livers and H&E staining 96 hours post-APAP administration (78 hours post-treatment) (left panel) (scale bars = 500 μm). Dotted lines indicate necrotic areas. Quantification of the necrotic area in three H&E-stained sections per mouse liver sample (right panel) (n = 4 mice per group). (m) Immunostaining for cleaved caspase-3 (cCasp3) (red) in the pericentral region of the liver, with nuclei counterstained with DAPI (blue) (scale bar = 100 μm). CV indicates the central vein. (n) Western blot analysis for cleaved caspase-3 (cCasp3) 96 h post-APAP administration (78 h post-LNP injection at indicated mRNA doses). Vinculin was used as an internal control. Statistical analysis was performed using one-way ANOVA: p < 0.01(**), p < 0.001(***), ns, not significant.

Finally, we investigated whether OSKM mRNA-LNP treatment could promote liver regeneration following AILI. Upon OSKM induction, the production of LPLCs, as indicated by Sox9 expression, began as early as 24 h (Fig. 5c). To achieve maximal OSKM expression at 24 h, OSKM mRNA-LNPs were administrated 18 h after APAP administration. The in vivo effect of OSKM-mRNA LNP on liver regeneration was then monitored (Fig. 6i). The typical sign of necrosis (Supplementary Fig. 6j) and a marked increase in serum AST and ALT levels (Supplementary Fig. 6k) were observed following administration of a sub-lethal dose of APAP (400 mg/kg). The survival rate of the OSKM mRNA-LNP-injected group was significantly higher compared to the Luc mRNA-LNP-treated group (Fig. 6j). OSKM mRNA-LNP-treated group exhibited comparable levels of serum AST and ALT to normal liver of healthy control at 96 hours post-APAP administration, whereas the mock group and Luc mRNA-LNP-treated group showed elevated levels of serum AST and ALT (Fig. 6k). Notably, the OSKM mRNA-LNP-treated mice exhibited a substantial reduction in necrotic area (Fig. 6l) and cell death in the pericentral region (Fig. 6m), corresponding to the decrease in caspase 3 activation in the 0.3 mg/kg OSKM mRNA-LNP-treated group (Fig. 6n). We validated that these therapeutic effects resulted from hepatic reprogramming following the liver-targeted delivery of OSKM mRNAs. The OSKM mRNA-LNP-treated mice exhibited a significant increase in the Sox9+ LPLC population, accompanied by noticeable proliferation, as evidenced by the Sox9 and Ki-67 levels at 48 h post-APAP administration (30 h after OSKM mRNA-LNP injection) (Supplementary Fig. 6l). Western blot analysis further confirmed the marked expression of Sox9 protein in these mice (Supplementary Fig. 6m). Moreover, we have conducted an extended study to investigate potential hepatic abnormalities beyond 96 hours post-injury following OSKM mRNA-LNP treatment. Histological analysis of liver harvested 30 days following OSKM mRNA-LNP injection (31 days post-injury) revealed no evidence of tumorigenesis (Supplementary Fig. 6n). No significant histological differences were observed among all groups, because acute liver injury induced by APAP had largely recovered by day 30 (Supplementary Fig. 6n). Masson’s trichrome (MT) staining and quantitative PCR analysis for *Col1a1* and *Acta2* demonstrated minimal fibrosis in the liver 30 days after injection (Supplementary Fig. 6o). Moreover, serum levels of AST, ALT, and albumin in mice 30 days following OSKM mRNA-LNP injection were comparable to those of the healthy control group (normal) (Supplementary Fig. 6p), demonstrating recovery of impaired hepatic function from liver injury in long-term monitoring. Consistent with the outcomes of OSKM transgene induction (Fig. 2 and Supplementary Fig. 2), the liver-targeted delivery of OSKM mRNA-LNP facilitated the partial reprogramming for hepatocyte dedifferentiation, presenting a promising therapeutic application for in vivo reprogramming in the liver.

## Discussion

Emerging evidence suggests that in vivo reprogramming with OSKM promotes both regeneration and rejuvenation^9–11,13,14^. Our previous study demonstrated that transient OSKM induction creates distinct populations of ‘injury-responsive cells’ in the intestine, suggesting that short-term OSKM induction can replicate the epigenetic changes associated with injury-induced reprogramming (or adaptive cellular reprogramming^2^)^13^. In the current study, we facilitated liver regeneration through the formation of LPLCs using short-course OSKM induction without injury. This approach not only accelerated regeneration following APAP injury but did so with minimal immune response (Fig. 4, 5) compared to the natural injury response. Interestingly, the induction of Sox9 and the subsequent activation of Stat3, triggered by IL-6 from activated Kupffer cells following APAP injury^5^, coincided with similar molecular changes induced by OSKM (Fig. 5). Notably, IL-6, a key cytokine in liver regeneration following hepatectomy^44^, is known to be produced by hepatocytes themselves in this context^45,46^. These findings raise the possibility that OSKM-mediated epigenetic reprogramming promotes autonomous IL-6 production by hepatocytes, thereby activating Stat3 signaling in a cell-intrinsic manner. This parallel suggests that LPLC formation may involve shared dedifferentiation mechanisms, similar to those previously demonstrated in the intestinal epithelium^13^.

The promising results of facilitated liver repair through OSKM-mediated cellular reprogramming underscore the potential for developing liver-specific transient OSKM induction strategies. While previous studies have utilized OSKM for tissue regeneration in various contexts^9,47^, the use of AAV vectors for long-term expression poses significant risks, including teratoma formation^48^ and premature mortality^29^, associated with intestinal and liver failure in a Dox-inducible reprogramming mouse model^28^. These findings highlight critical concerns regarding the potential hazards of sustained gene expression, even over relatively short durations.

Alternatively, LNP-based mRNA delivery systems offer significant advantages due to their capacity for transient expression of target genes and precise control over the duration of gene expression by adjusting the dosage and frequency of administration. This approach mitigates the toxicity associated with prolonged gene expression and ensures safety^41^. Protein expression in the liver following mRNA delivery via LNPs was achieved within 24 h and markedly diminished by 48 h after a single intravenous administration (Fig. 6c). Upon intravenous administration, most of the LNPs are adsorbed by Apolipoprotein E (ApoE) serum proteins, leading to predominant delivery to hepatocytes^49,50^. The passive targeting ability of LNPs to the liver suggests that the liver is an optimal target organ for in vivo partial reprogramming.

Importantly, we enhanced liver-specific OSKM mRNA delivery using LNPs conjugated with galactose moieties to specifically target hepatocytes. Successful delivery of OSKM mRNAs to the liver using LNPs was achieved without a significant increase in serum AST and ALT levels, indicating the lack of hepatotoxicity. We demonstrated that the OSKM mRNA-LNP platform could replicate the injury-induced dedifferentiation process in the liver and promote liver regeneration post-AILI. OSKM mRNA-LNP induced hepatocyte dedifferentiation, LPLC production, and liver regeneration after sub-lethal AILI. The transient expression of reprogramming factors following OSKM mRNA-LNP administration suggests a controllable partial reprogramming strategy, mitigating the detrimental effects of sustained in vivo reprogramming. Consistent with previous findings in the reprogramming mouse model^12^, our results support the potential of OSKM mRNA delivery via LNPs as a safe and effective method for hepatic OSKM induction aimed at liver regeneration.

The LNP-mRNA platform holds significant potential for developing therapeutic strategies to address various diseases and pathological conditions beyond acute liver injury. Given the recent development of organ-specific LNP-based mRNA delivery systems for extrahepatic tissues (e.g., lung and spleen)^51–53^, this approach offers promising therapeutic options for treating refractory diseases that require tissue regeneration and rejuvenation in non-hepatic organs through in vivo partial reprogramming.

## Materials and Methods

### Mice

Col1A1^TetO-OSKM^;ROSA26^rtTA^ mice (iOSKM mice) were obtained from the Jackson Laboratory (no. 011004). ROSA26^rtTA^ mice were generated from Col1A1^TetO-OSKM^;ROSA26^rtTA^ mice. Mice used for all experiments were 8-12 weeks of age and housed in conventional cage with 12h-12h light-dark cycle and access to food and water ad libitum, under pathogen-free conditions at the Seoul National University. Most experiments were conducted with male mice unless described in the figure legends. For OSKM induction, mice were treated with Doxycycline hyclate (Sigma, D9891) 0.15 mg/ml in drinking water containing 5% sucrose (Samchun Chemicals, 50010) for one to four days. For BrdU pulse-chase experiment, BrdU (Sigma, B5002) 150 mg/kg in PBS was injected intraperitoneally 2 hours before euthanasia. For acetaminophen induce liver injury experiment, Acetaminophen (Sigma, A7085) was intraperitoneally injected after fasting for 12h (10:00 pm ∼ 10:00 am), with or without doxycycline treatment. Acetaminophen was solved in normal saline (20 mg/ml) and kept warm to avoid precipitation. These animal experiments were conducted under the permission of Seoul National University Institutional Animal Care and Use Committee (permission number: SNU-210326-5, SNU-240125-1). For the OSKM mRNA-LNP experiment in acute liver injury mouse model, LNP-mRNAs were intravenously injected to wild-type C57BL/6 mice (male, 8-12 weeks old) (Orient Bio, Seongnam, Korea) via retro-orbital venous sinus injection 18 hours post-APAP administration. LNP-mRNAs were diluted to the desired concentration in phosphate buffered saline (PBS; ThermoFisher Scientific, Waltham, MA, USA) prior to each experiment, and 100 μl of the diluted LNPs loading firefly luciferase or OSKM mRNA was intravenously injected at an mRNA dose of 0.3 mg/kg (or 0.6 mg/kg). These animal experiments were approved by the Institutional Animal Care and Use Committee (IACUC) of Yonsei University (permission number: IACUC-A-202301-1618-02).

### Hepatocyte isolation

Mice were anaesthetized with Avertin (250 mg/kg, IP injection) before surgical procedure. After anaesthetization, the hepatocyte isolated with well-established two-step collagenase perfusion ^54^ with some adjustments. Briefly, 24 G syringe connected to perfusion tube and peristaltic pump was inserted into the portal vein and vena cava was cut after cannulation for allowing blood flow. 10 ml of EDTA buffer was perfused to wash out the blood from liver. Continuously, 20 ml of collagenase buffer (1 mg/ml) was perfused for digestion. While the collagenase buffer perfusion, several times of clamping the vena cava with forceps for 5∼10 s was conducted for efficient perfusion to all liver vasculatures. Perfused liver was dissected and moved to Petri dish with 20ml of hepatocyte culture media and ruptured with forceps to release hepatic cells. All hepatic cells were filtered through 70μm cell strainer. Cells were centrifuged at 50 g for 2 min at 4 °C and the supernatant were discarded to eliminate non-parenchymal cells. Hepatocyte-enriched pellet was resuspended in HBSS and centrifuged at 50 g for 2 min at 4 °C for washing. To enrich the live hepatocyte, 9 ml of Percoll (Sigma, P1644) mixed with 1 ml of 10X PBS was added to HBSS contain hepatocytes. Cells were centrifuged at 200 g for 5 min at 4 °C. Live hepatocytes were kept for further analysis or culture.

### In vitro culture of primary hepatocyte

For in vitro culture of primary hepatocytes, isolated primary hepatocytes were counted by trypan blue staining using hemacytometer and 1×10^6^ cells were plated in collagen-coated 60π culture plates. For maintenance of primary hepatocytes, cells were grown in Dulbecco’s modified Eagle’s medium (DMEM) supplemented with 10% FBS and 1% penicillin/streptomycin. Culture media was changed 3 hours after plating and refreshed every 24 hours. For OSKM induction, primary hepatocytes were treated with doxycycline (0.5 or 1 μg/ml) for 48 hours.

### Immunohistochemistry and immunofluorescence

For paraffin embedded sections, liver samples were fixed in 4% PFA at 4°C overnight and embedded in paraffin blocks next day. For frozen sections, each liver samples were fixed in 4% PFA at 4°C overnight and immersed in 30% sucrose at 4°C overnight. Prepared liver samples cryopreserved in Tissue-Tek optimal cutting temperature (OCT) compound (Sakura Finetek).

For immunohistochemistry, 4 μm paraffin sections were treated with 0.3 % H_2_O_2_ in methanol for blocking endogenous peroxidase activity. For antigen retrieval, slides were kept in tris-EDTA buffer (pH 9.0) at 95°C for 20 min. Retrieved sections were washed with TBST (0.1 % Triton X-100 in TBS) for 10 min and blocked by incubation with 10% normal goat serum in TBST for 1 hour. Then, slides were incubated with primary antibody at 4 °C overnight or at room temperature for 1 hour. Subsequently, the slides were incubated with a HRP-conjugated secondary antibody at room temperature for 2 hours, followed by 2- to 6-min incubation with 3,3′-diaminobenzidine substrates (Vector Laboratories, SK-4100). Hematoxylin (Sigma, GHS3) staining was conducted for nuclear counterstaining. Slides were covered with slide glass using MOWIOL mounting solution. Samples were visualized with a brightfield microscope (Leica DM500) or Automated multimodal tissue analysis system (PerkinElmer, Vectra). For immunofluorescence, 8 μm frozen sections were incubated with primary antibodies in 4% bovine serum albumin (BSA) in TBST at 4 °C overnight and subsequently incubated with fluorescent secondary antibodies at room temperature for 2 hours. 4′,6-Diamidino-2-phenylindole (DAPI) staining was conducted for nuclear counterstaining. Slides were covered with slide glass using MOWIOL mounting solution. Samples were visualized with a fluorescence microscope (Olympus BX53 or Leica THUNDER Imager). Following antibodies were used for immunostaining : Sox9 (Millipore, AB5535, 1:500), Hnf4α (Abcam, ab41898, 1:400), Ki-67 (Abcam, ab16667, 1:400), BrdU (Novus Biologicals, NBP2-14890, 1:200), Oct4 (BD, 611203, 1:500), Sox2 (Millipore, AB5603, 1:400), F4/80 (Cell Signaling Technology, 70076, 1:400) and cleaved Caspase-3 (Cell Signaling Technology, 9664, 1:400).

### Histological analysis

Liver samples were harvested and fixed in 10% formalin solution at 4 °C overnight. The fixed tissues were embedded in paraffin blocks, sectioned into slices of 5 μm thickness, and stained with Hematoxylin (H9627, Sigma-Aldrich) and Eosin Y (000E0614, Samchun Chemicals, Seoul, Korea) (H&E) or Masson’s trichrome (B8563, Sigma-Aldrich). For staining of glycogen, sections were stained Periodic Acid Schiff Stain Kit (ScyTek Laboratories, PAS-2-IFU) following the manufacturer’s protocol. Dehydrated samples covered with slide glass using Canada balsam (Sigma-Aldrich). Samples were visualized with a microscope (Leica DM500). Images were obtained with slide scanner (VS120-S5-W, Olympus, Tokyo, Japan) or brightfield microscope (Leica DM500).

### Blood chemistry analysis

Blood samples were collected from mice for serum biochemical analyses. The blood samples were allowed to clot at room temperature for at least 30 minutes and then centrifuged at 2000 g for 10 minutes at 4 °C to obtain serum. Biochemical analyses for alanine aminotransferase (ALT), aspartate aminotransferase (AST), and albumin (ALB) were conducted as indicators of liver function using the DRI-CHEM 4000i (Fuji Film, Tokyo, Japan).

### RNA isolation and RT-qPCR

Total RNA of isolated hepatocytes or whole liver was extracted using Easy-BLUE total RNA extraction kit (iNtRON Biotechnology, 17061). cDNA synthesis with total RNA samples was performed using the PrimeScript RT reagent kit (TAKARA, RR036A) according to the manufacturer’s protocol. RT-qPCR was performed with QuantStuido 3 (ThermoFisher Scientific) using TB-green (TAKARA, RR420) following the manufacturer’s instructions.

### Immunoblotting

Hepatocytes or whole liver samples were lysed in RIPA buffer containing 1% protease inhibitor cocktail and 0.1% sodium orthovanadate on ice for 1 hour with periodic vertexing. Lysates were clarified by centrifugation. Total protein concentration was determined using the BCA protein assay kit (ThermoFisher Scientific, #23225). Approximately 15-20 μg of total protein were separated by SDS-PAGE on gels with varying acrylamide concentrations (7.5%, 10%, 12%, 15%). Separated proteins were transferred to PVDF membranes. The membranes were blocked with 5% skim milk in TBST (Tris-buffered saline with 0.1% Tween-20) for 1 hour at room temperature and washed twice with TBST for 10 minutes each. Membranes were then incubated overnight at 4°C with primary antibody diluted in TBST containing 0.1% sodium azide. Following incubation, membranes were washed three times with TBST for 10 minutes each. Membranes were subsequently incubated with HRP-conjugated secondary antibody (Jackson Immunoresearch Laboratories) in TBST for 1 hour at room temperature. Finally, membranes were washed three times with TBST for 15 minutes each. Immunoreactive bands were detected using the WEST-Queen kit (iNtRON Biotechnology, #16026) on a ChemiDoc system.

### Flow cytometry for assessment of hepatocyte ploidy

Freshly isolated hepatocytes were fixed with 75% ethanol in PBS at room temperature for 15 min. Fixed hepatocytes were treated with 1% RNase A in PBS at room temperature for 30 min with periodic inverting. RNase treated hepatocytes were stained PI solution (1 mg/ml in DW) and analyzed with a flow cytometer (BD FACSCelesta). The data were analyzed with FlowJo software.

### TUNEL assay

TUNEL assay was performed using an in situ apoptosis detection kit (RnD Systems, 4810-30-K). To assessment of apoptotic cell death after acute liver injury, the apoptotic area stained with 3,3′-diaminobenzidine (DAB) were evaluated in liver sections. Hematoxylin (Sigma-Aldrich, GHS3) staining was conducted for nuclear counterstaining. Slides were covered with slide glass using MOWIOL mounting solution. Samples were visualized with a brightfield microscope (Leica DM500).

### Materials used for the formulation of LNPs

C12-SPM lipid was synthesized from Medigen (Daejeon, Korea) through conjugate addition of alkyl epoxides to polyamines via ring-opening reactions, as in the previous study ^42^. DOPE (#850725), C16-PEG2000 ceramide (#880180), DMG-PEG2000 (#880151), and C16 Galactosyl(α) ceramide (#860431) were obtained from Avanti Polar Lipids (Alabaster, AL, USA). DSPC (P1138) and cholesterol (C8667) were purchased from Sigma-Aldrich (Saint Louis, MO, USA). DLin-MC3-DMA was obtained from Med Chem Express (HY-112251, Monmouth Junction, NJ, USA). Firefly luciferase mRNA was purchased from TriLink BioTechnologies (L-7202, San Diego, CA, USA). Oct4, Sox2, Klf4, and c-Myc mRNA were produced through the in vitro transcription (IVT) method.

### In vitro transcription for mRNA preparation

Oct4, Sox2, Klf4, and c-Myc plasmids, containing poly T tail structure in the antisense strand, were linearized using the NotI restriction endonuclease (R001S, New England Biolabs, Ipswich, MA, USA), and transcribed into mRNAs using the MEGAscript T7 Transcription Kit (AM1334, Invitrogen, Waltham, MA, USA). The mRNAs were capped via the co-transcriptional capping method, using 3’-O-Me-m7G(5’)ppp(5’)G RNA Cap Structure Analog (S1411L, New England Biolabs). To generate nucleoside-modified mRNAs, N1-Methylpseudourindine-5’-Triphosphate (N-1081, TriLink BioTechnologies) was used instead of UTP. In vitro transcribed mRNAs were analyzed by 1% agarose gel electrophoresis for the verification of sequence size and purity, and were stored at -80 °C before use.

### Formulation and characterization of LNP-mRNA

As the organic phase for the formulation of C12-SPM-GAL LNP, ionizable lipid C12-SPM, DOPE, cholesterol, C16-PEG2000 ceramide, and C16 Galactosyl(α) ceramide were dissolved in ethanol at a molar ratio of 20:26.5:50.5:1.5:1.5. For DLin-MC3-DMA LNP, DLin-MC3-DMA, DSPC, cholesterol, DMG-PEG2000 were solubilized in ethanol at a molar ratio of 50:10:38.5:1.5. mRNAs were dissolved in 10 mM citrate buffer (pH3) (854, Sigma-Aldrich) as the aqueous phase. The aqueous and organic phases were mixed at a volume ratio of 3:1 using a microfluidic mixing device (Ignite, Precision NanoSystems, Vancouver, Canada). The formulated LNPs were incubated at room temperature for 15 minutes and dialyzed against phosphate buffered saline (PBS; 10010-023, ThermoFisher Scientific) using 3.5 kDa MWCO dialysis cassettes (66330, ThermoFisher Scientific) overnight at 4 °C. LNPs were then concentrated using ultracentrifugal filters (UFC901096, Merck Milipore, Burlington, MA, USA), filtered through 0.22 μm filter, and stored at 4 °C. mRNA encapsulation efficiency was analyzed using Quant-iT RiboGreen RNA assay kit (R11490, Invitrogen). The hydrodynamic diameter, zeta potential, and polydispersity index (PDI) of the LNPs were measured by dynamic light scattering (DLS) using Zetasizer Nano ZSP (Malvern Instruments, Malvern, United Kingdom).

### In vitro transfection of LNP-mRNA

Primary mouse embryonic fibroblasts (MEFs) were isolated from ICR mouse embryos at E13.5 days (Orient Bio) as previously described ^55^. Protocols for the animal experiment were approved by the Institutional Animal Care and Use Committee (IACUC) of Yonsei University (permission number: IACUC-202201-1407-08). Isolated MEFs were cultured on 0.2% gelatin-coated plates in Dulbecco’s modified Eagle’s medium (DMEM; 11995-065, ThermoFisher Scientific) supplemented with 10% (v/v) fetal bovine serum (FBS; 26140079, ThermoFisher Scientific). One day after seeding onto 12-well plates, LNP-mRNA solutions were diluted in reduced-serum medium (Opti-MEM; 31985070, ThermoFisher Scientific) at an mRNA concentration of 10 μg/ml, and 100 μl of the diluted LNP-mRNA solutions were added dropwise (1 μg mRNA per 100,000 cells) into the cell medium and incubated at 37 °C for 24 hours. One day after the transfection, cells were lysed for protein extraction or analyzed for cell viability and immunocytochemistry.

### In vivo bioluminescence imaging

C57BL/6 mice (8∼12 weeks old) (Orient Bio) were intravenously injected with LNPs loaded with luciferase mRNA (0.3 mg/kg, n=3∼4 per group). After 6, 12, 24, 48, and 96 hours post-LNP injection, these mice were anesthetized with 2% isoflurane (657801261, Hana Pharm, Seoul, Korea) and intraperitoneally injected with D-Luciferin (P1043, Promega, Madison, WI, USA) at a dose of 150 mg/kg. Bioluminescence was detected 10 minutes after luciferin administration using an IVIS Spectrum imaging system (PerkinElmer, Waltham, MA, USA), while maintaining 2% isoflurane in the imaging chamber, and quantified using Living Image software (PerkinElmer). For ex vivo organ bioluminescence analysis at 6 hours post-LNP injection, five major organs (heart, lungs, liver, spleen, and kidney) were collected 10 minutes after D-luciferin administration, and bioluminescence signals were detected using an IVIS Spectrum imaging system. These animal experiments were approved by the Institutional Animal Care and Use Committee (IACUC) of Yonsei University Health System (permission number: 2022-0327).

### Cell viability measurement

To evaluate cell viability after the transfection of LNP-mRNA, MEFs treated with each LNP-mRNA for 24 hours were incubated with methylthiazolyldiphenyl-tetrazolium bromide (MTT) solution (M2128, Sigma-Aldrich) for 3 hours at 37 °C. The resulting formazan in each well was dissolved in dimethyl sulfoxide (DMSO; 046-21981, Wako, Osaka, Japan), and the absorbance of the dissolved solution at 560 nm was measured using a microplate reader (Infinite M200, TECAN, Männedorf, Switzerland).

### Immunocytochemistry

Cells were fixed with 10% formalin solution (HT501128, Sigma-Aldrich) for 10 minutes at room temperature and washed with PBS three times. Fixed cells were permeabilized with 0.2% (v/v) Triton X-100 (X100, Sigma-Aldrich) for 15 minutes, and then blocked with 4% (w/v) bovine serum albumin (BSA; 02160069, MP Biomedicals, Santa Ana, CA, USA) for an hour. The cells were subsequently incubated with the following primary antibodies overnight at 4 °C: mouse anti-Oct3/4 (1:200, sc-5279, Santa Cruz Biotechnology, Dallas, TX, USA) and mouse anti-Sox2 (1:200, sc-365823, Santa Cruz Biotechnology). After washing with PBS three times, the samples were then incubated with the following secondary antibodies for an hour at room temperature: Alexa Fluor 488-or Alex Fluor 594-conjugated anti-mouse IgG (1:200, A11001, A11005, ThermoFisher Scientific). Nuclei were counterstained with 4’,6-diamidino-2-phenylindole (DAPI; A2412, TCI America, Portland, OR, USA). Images were obtained with confocal microscopy (LSM 880, Zeiss, Oberkochen, Germany).

### Bulk RNA-seq data analysis

RNA-seq data of liver post-AILI were obtained from public dataset (https://doi.org/10.5281/zenodo.6035873). This dataset contains UMI counts for a total of 17,999 genes across 21 liver samples. To analyze differential gene expression between AILI and control groups, we employed Wald tests, which were implemented in the R package DEseq2. Differentially expressed genes (DEGs) were selected based on adjusted p-value less than 0.01 and an absolute log fold-change greater than 1 as the cut-offs. Gene set enrichment analysis (GSEA) was conducted to yield pathways significantly overrepresented at the top or the bottom of ranked gene lists acquired from the DEG analysis between time-course groups. The GSEA was performed using the R package fgsea. The normalized enrichment score (NES) was calculated and applied to generate radar plot. Additionally, soft clustering was employed to group genes based on their expression patterns across the time-course groups. Soft clustering was performed using the R package Mfuzz. The optimal fuzzifier value were estimated with ‘mestimate’ function.

### snRNA-seq data analysis - Quality control

Before quality control (QC), our dataset contained 47,266 cells and 32,285 genes. QC was performed within individual samples by calculating trimmed mean (t_mean) and standard deviation (t_std) with 10% of trimming. Filtered categories were sample’s total read number (total counts), total gene number (gene counts), hemoglobin gene number (hb counts), and mitochondrial gene number (mt counts).

Cells with ’total counts’ and ’gene counts’ below (t_mean - 1*t_std) or above (t_mean + 4*t_std) were filtered out. Cells with ’hb counts’ above 4*t_std and ’mt counts’ above 1*t_std were also filtered out. Additionally, genes that were not expressed in more than 5 cells were removed. This left us 25,210 cells and 20,349 genes. Finally, in addition to scrublet results ^56^, we manually curated low-quality cells or doublets, resulting in final cell number of 22,641. Read counts data were normalized with median counts per cell and log1p-normalized (base = e).

### snRNA-seq data analysis - Batch correction and embedding

We performed scVI and scANVI ^57^ for data embedding using references from public datasets ^58–60^. In merged reference data, we identified highly variable genes (HVG). Firstly, we ranked HVG across all genes, and selected genes that expressed more than 80% of each sample and more than 12 samples. Then we recalculated HVG and chose the top 6,000 genes. Likewise, we identified HVG on our snRNA-seq data. With the same method, we first selected genes that expressed over 90% in each sample and over 5 samples. We got an intersection of HVG sets from datasets, 3,372 genes in total. With the HVG and reference dataset, we performed scVI to calculate latent space. Batch correction was then performed using scANVI on the latent space derived from scVI. That latent space was used for embedding and reference mapping to our data. Additionally, we performed Scanorama for further batch correction ^61^. Scanorama latent space was used for compute knn neighbors and generating UMAP space. Lastly, cluster annotation was decided by expression of well-known marker genes and reference mapping results.

### snRNA-seq data analysis - Metacell selection and trajectory analysis

For trajectory analysis, we used the SEACell package to select metacells from hepatocyte cluster ^62^. Each metacell represented ∼75 cells. We used scFates package for trajectory calculation ^63^. Within each trajectory, we recalculated HVG in our metacell data. With the HVG, we conducted PCA and batch correction on the metacell data. Next, we calculated diffusion map derived from corrected latent space. The result further used as input for computing knn neighbor, UMAP space, and trajectory.

### GSEA (Gene-Set Enrichment Analysis)

GSEA was performed using GSEApy package ^64^. Gene lists for GSEA input was DEG results of scanpy package. Gene sets libraries used for GSEA are ’BioCarta_2016’, ’GO_Biological_Process_2023’,’KEGG_2019_Mouse’,’MSigDB_Hallmark_2020’,’Reactome_ 2022’,’WikiPathways_2019_Mouse’ from GSEApy libraries.

### Statistical analysis

Statistical analysis between two groups were performed using Student’s *t*-test, and in case of three or more groups with single variable, one-way analysis of variance (ANOVA) followed by Bonferroni comparison test was conducted. Two-way ANOVA was performed for groups with two or more variables. Dunnett’s multiple comparison test was performed in case of comparing experimental groups with single control group. Significance was set as P < 0.05 (*), P < 0.01 (**), P < 0.001 (***), P < 0.0001 (****). Error bars represent mean ± s.d.

## Data availability

Single-nucleus RNA sequencing data generated in this study are available in NCBI Gene Expression Omnibus (GEO) under the accession number GSE274988, https://www.ncbi.nlm.nih.gov/geo/query/acc.cgi?acc=GSE274988.

## Acknowledgements

This work was supported by the National Research Foundation of Korea (NRF) grant, funded by the Korean government through the Ministry of Science and ICT (MSIT) (RS-2024-00466703, RS-2023-00218543 and RS-2024-00432867 for HJ.C, and RS-2024-00432653 for SW.C). Global - Learning & Academic research institution for Master’s·PhD students, and Postdocs(LAMP) Program of the NRF grant funded by the Ministry of Education (No. RS-2024-00442483) for SH.H.

## Author contributions

HJ.C conceived the overall study design, led the experiments, and wrote the manuscript. S.H provided single-cell RNAseq data analysis and contributed to manuscript writing. SW.C led the LNP experiments and participated in manuscript writing. BK.J and YS.S conducted experiments, engaged in critical discussions of the results, and wrote the initial draft of the manuscript. W.S performed scRNAseq analysis. HJ.E, YJ.L, and J.K conducted animal experiments and sample preparation. All authors contributed to writing and revising the manuscript and approved the final version.

---

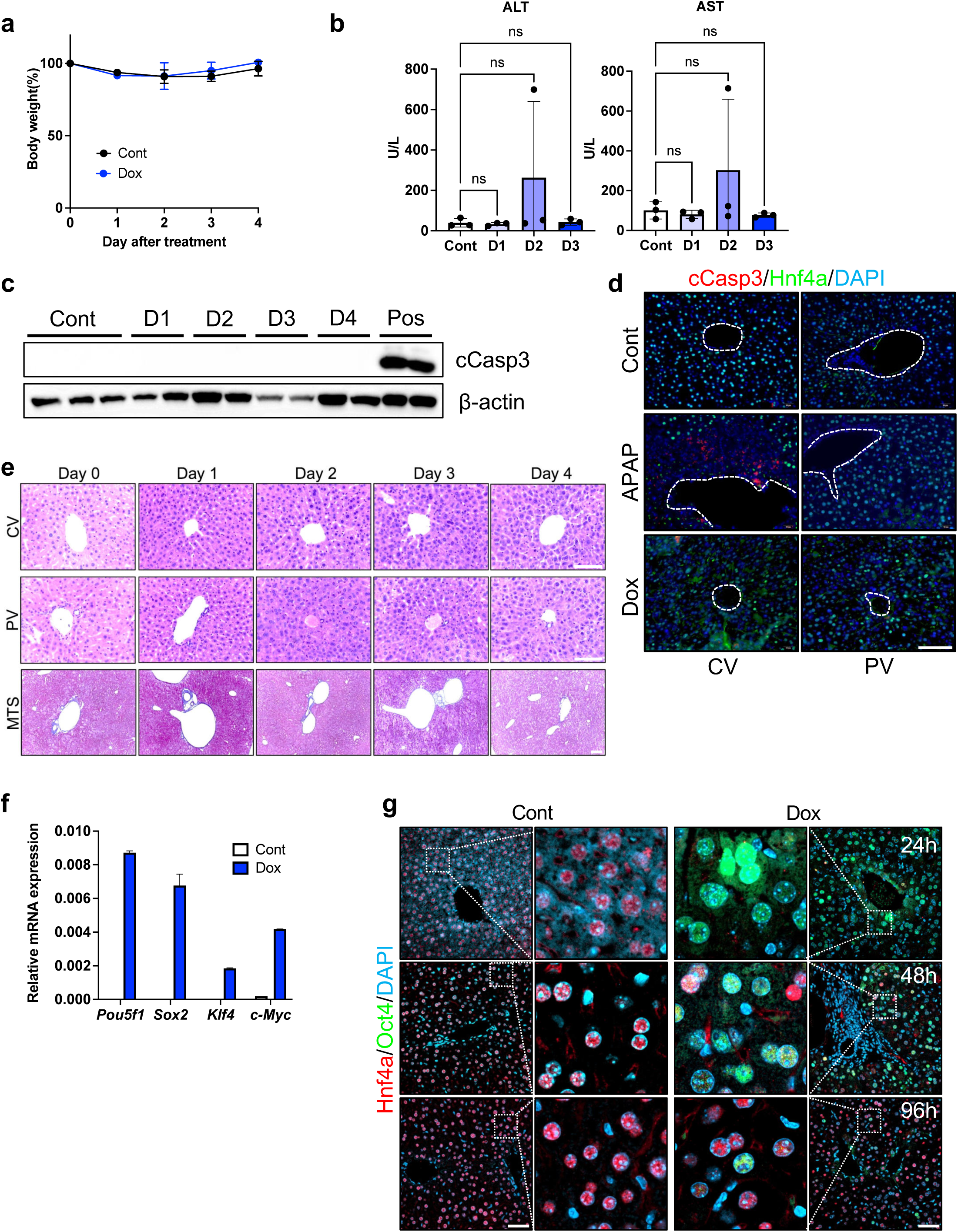
Supplementary Figure 1

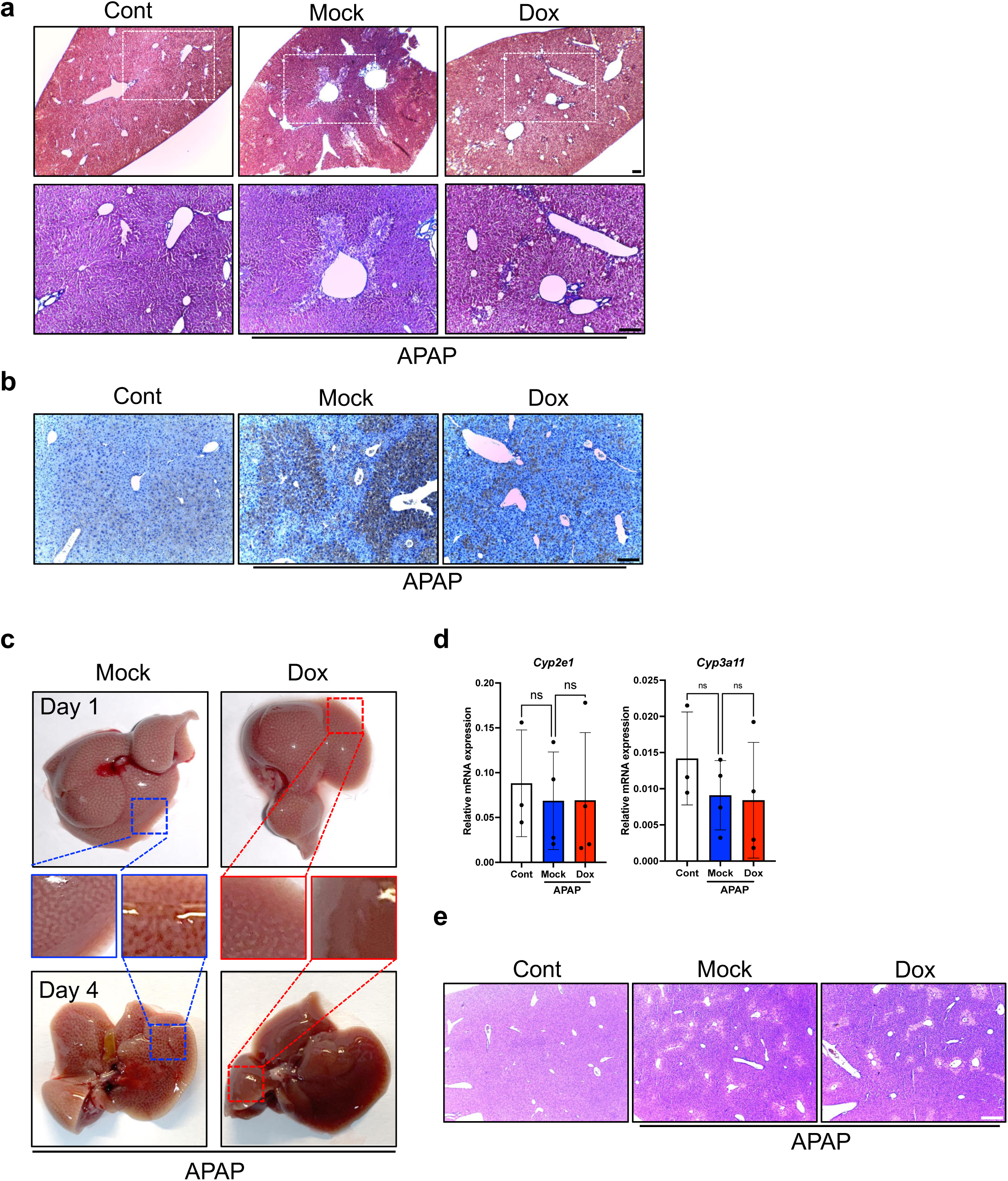
Supplementary Figure 2

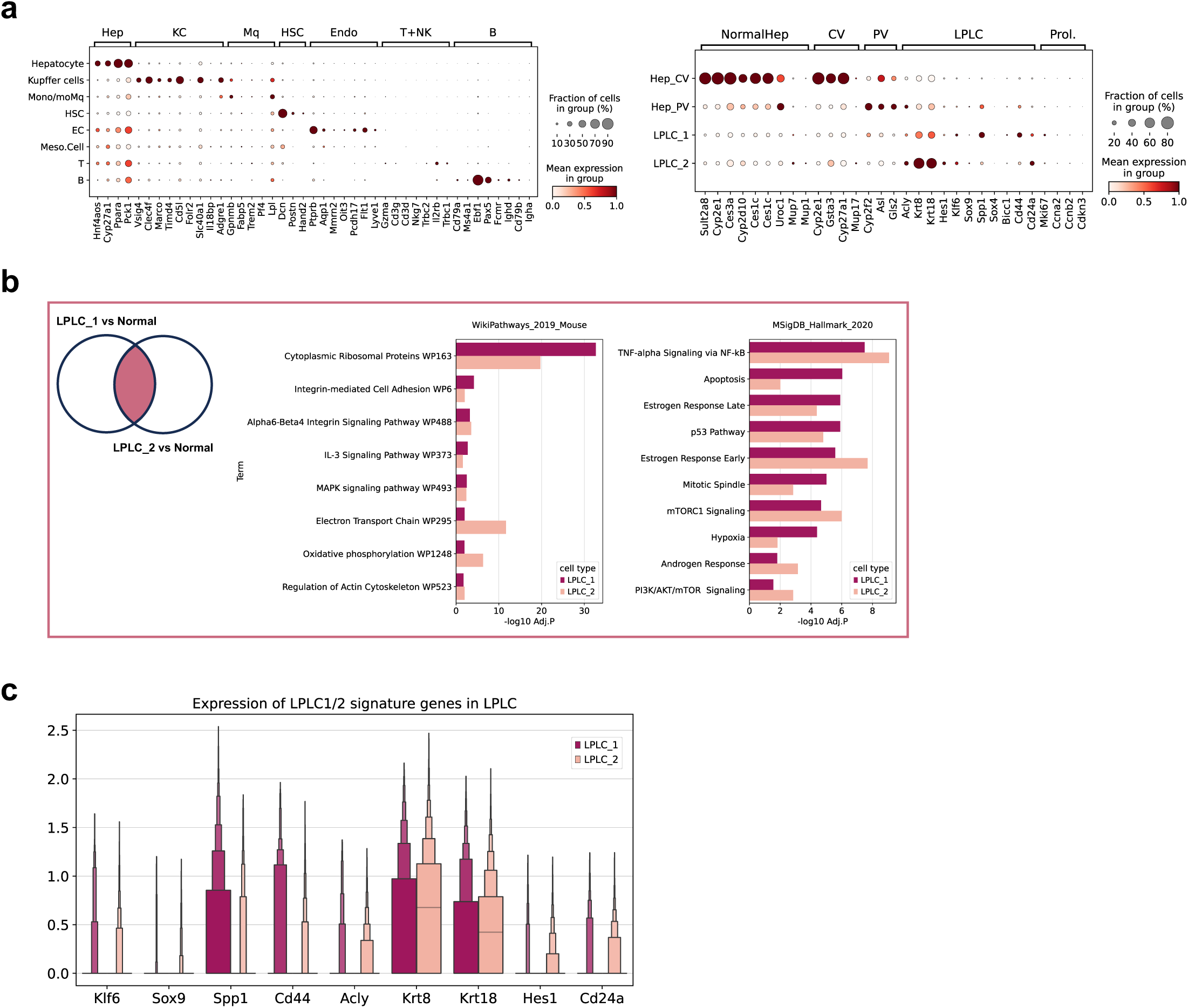
Supplementary Figure 3

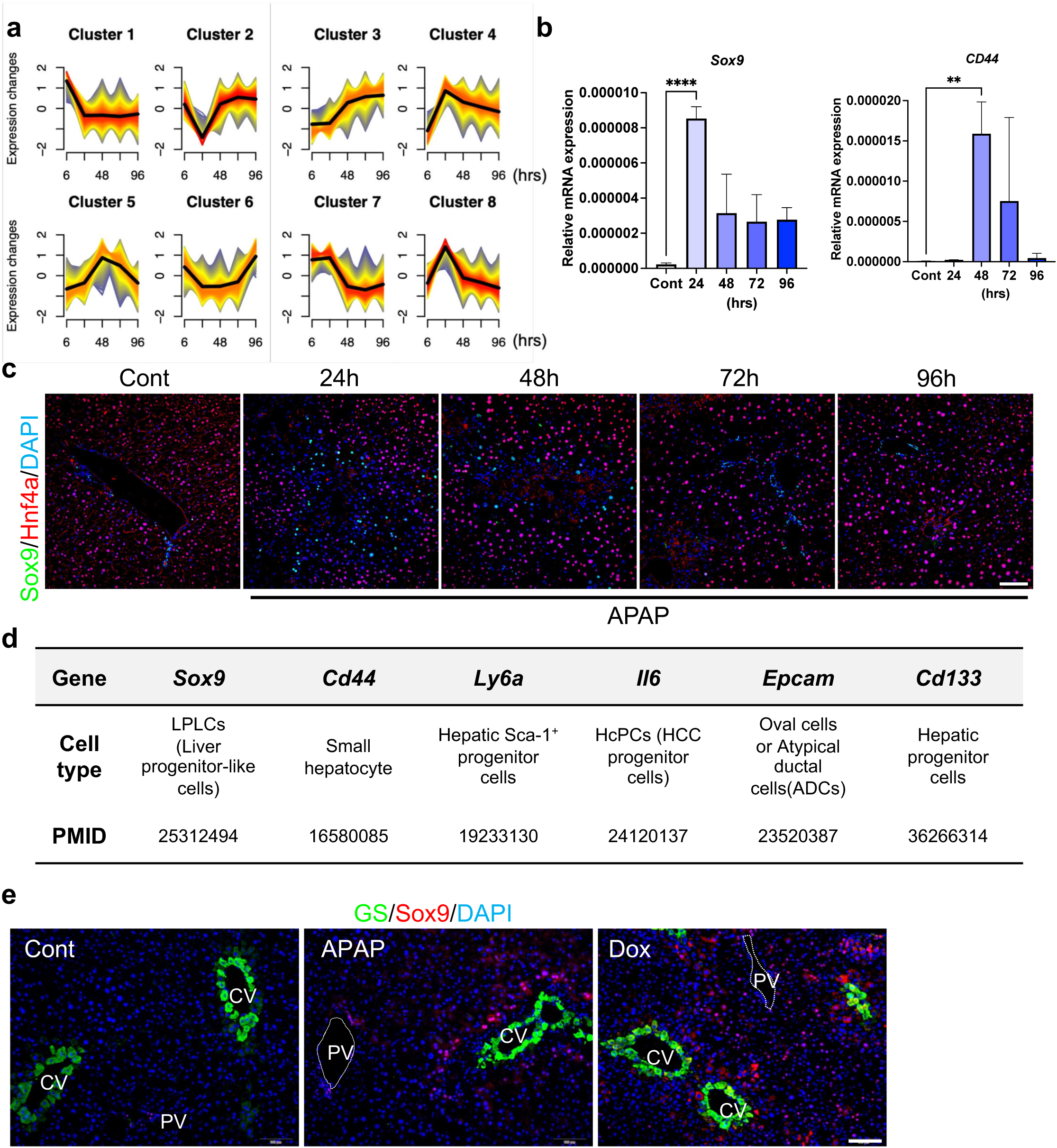
Supplementary Figure 4

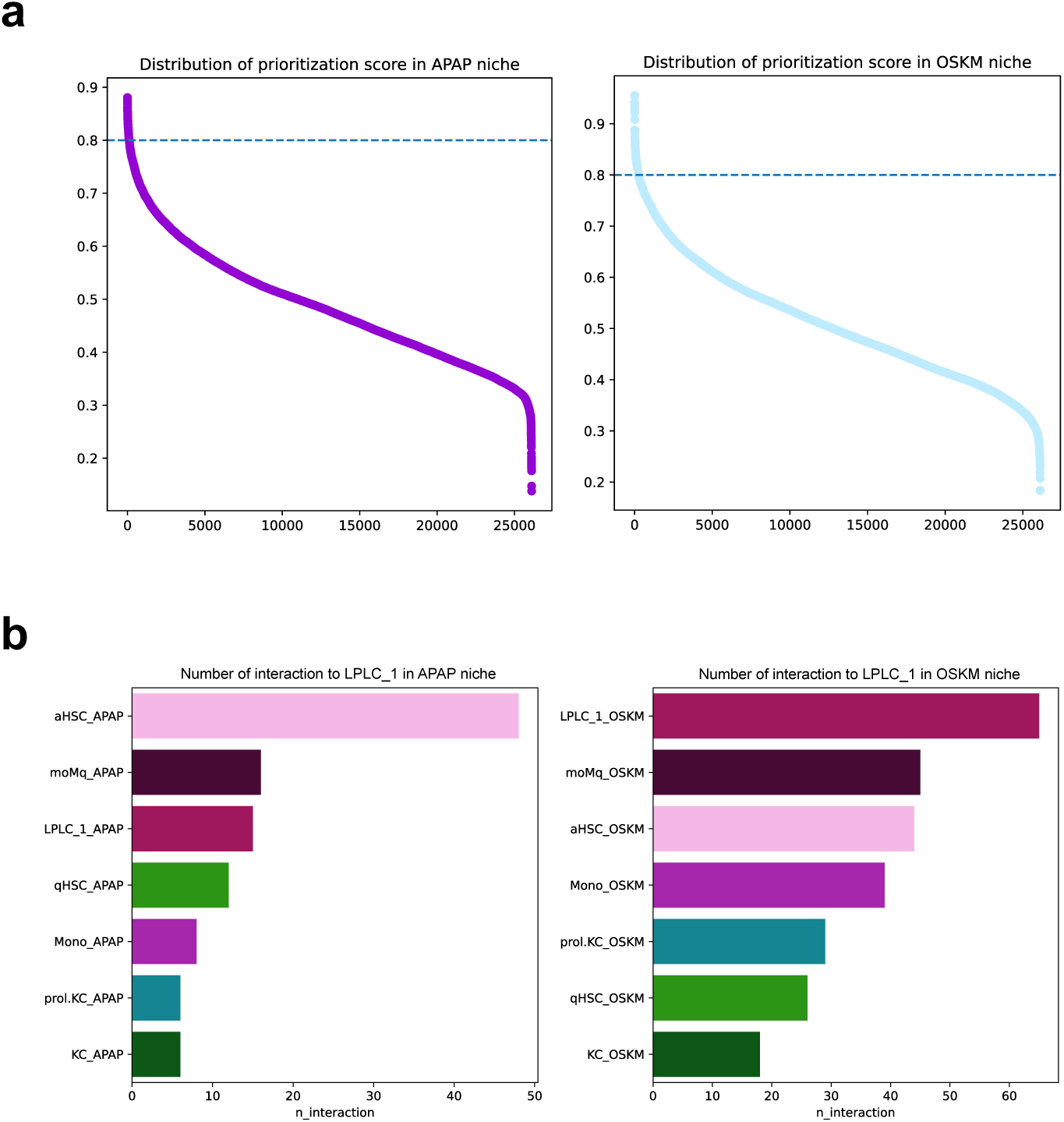
Supplementary Figure 5

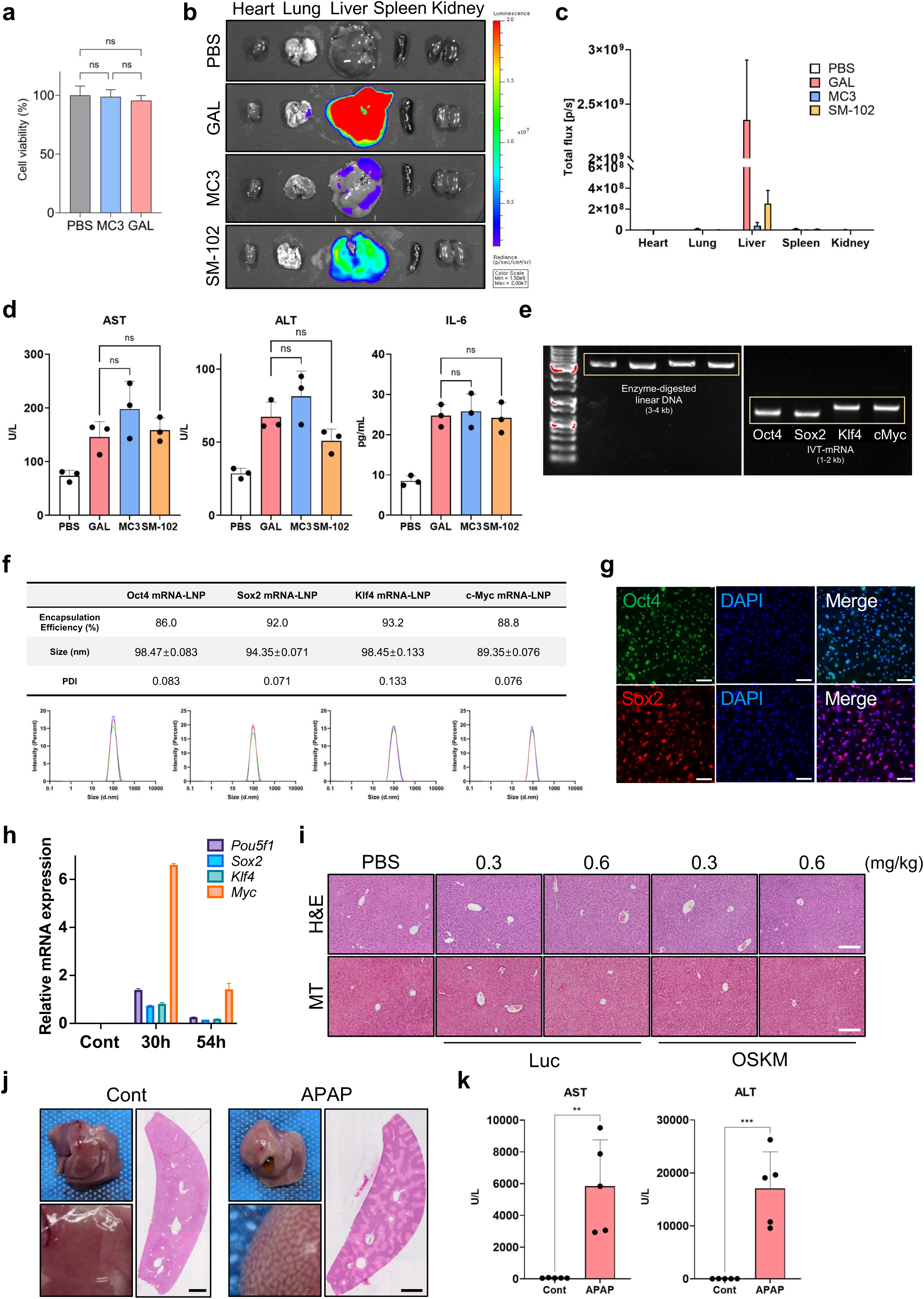
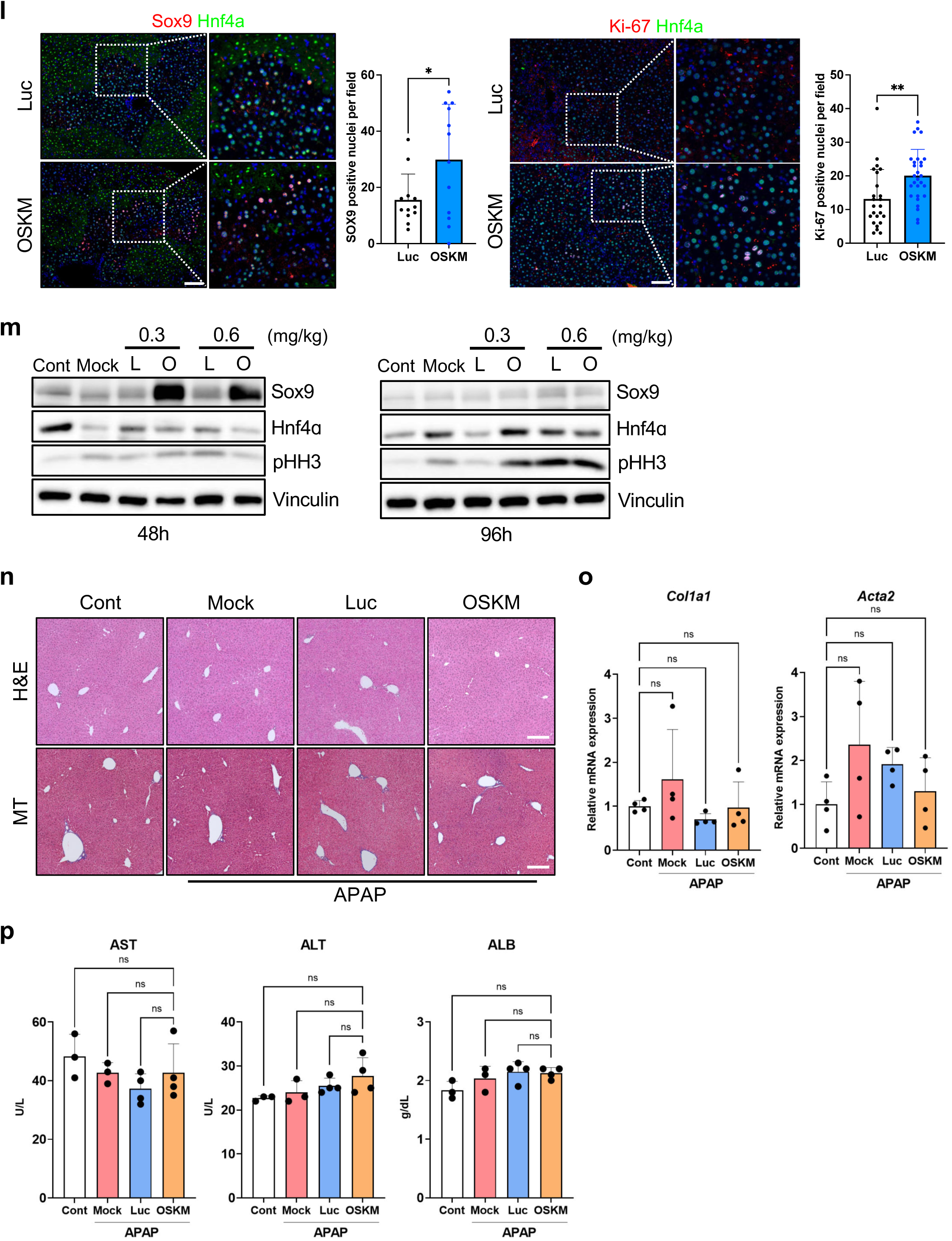
Supplementary Figure 6

