## Supplementary data for "Enhanced liver regeneration via targeted mRNA delivery for partial in vivo reprogramming"

**Running title**: Controlled mRNA delivery for in vivo reprogramming

Beom-Ki Jo[1#], Young Seok Song[3#], Woohyun Song[6,7#], Hee-Ji Eom[1], Yon Jae Lee[3], Jumee Kim[1], Seunghee Hong[6,7*] Seung-Woo Cho[3,4,5*], and Hyuk-Jin Cha[1,2*]

[1]College of Pharmacy, Seoul National University, Seoul, Republic of Korea, [2]Research Institute of Pharmaceutical Sciences, Seoul National University, Seoul, Republic of Korea, [3]Department of Biotechnology, College of Life Science and Biotechnology, Yonsei University, Seoul, Republic of Korea, [4]Cellartgen, Seoul, Republic of Korea, [5]Center for Nanomedicine, Institute for Basic Science (IBS), Seoul, Republic of Korea. [6]Department of Biochemistry, College of Life Science and Biotechnology, Yonsei University, Seoul, Republic of Korea, [7]Brain Korea 21 (BK21) FOUR Program, Yonsei Education & Research Center for Biosystems, Yonsei University, Seoul, Republic of Korea

[#]These authors contributed equally to this work.

*To whom correspondence should be addressed to

Prof. Hyuk-Jin Cha, Ph.D.

College of Pharmacy, Seoul National University

1 Gwanak-ro, Gwanak-gu, Seoul 08826, Republic of Korea

Prof. Seunghee Hong, Ph.D.

Department of Biochemistry, College of Life Science and Biotechnology, Yonsei University

50 Yonsei-ro, Seodaemun-gu, Seoul 03722, Republic of Korea

Prof. Seung-Woo Cho, Ph.D.

Department of Biotechnology, College of Life Science and Biotechnology, Yonsei University

50 Yonsei-ro, Seodaemun-gu, Seoul 03722, Republic of Korea

### Supplementary Figures

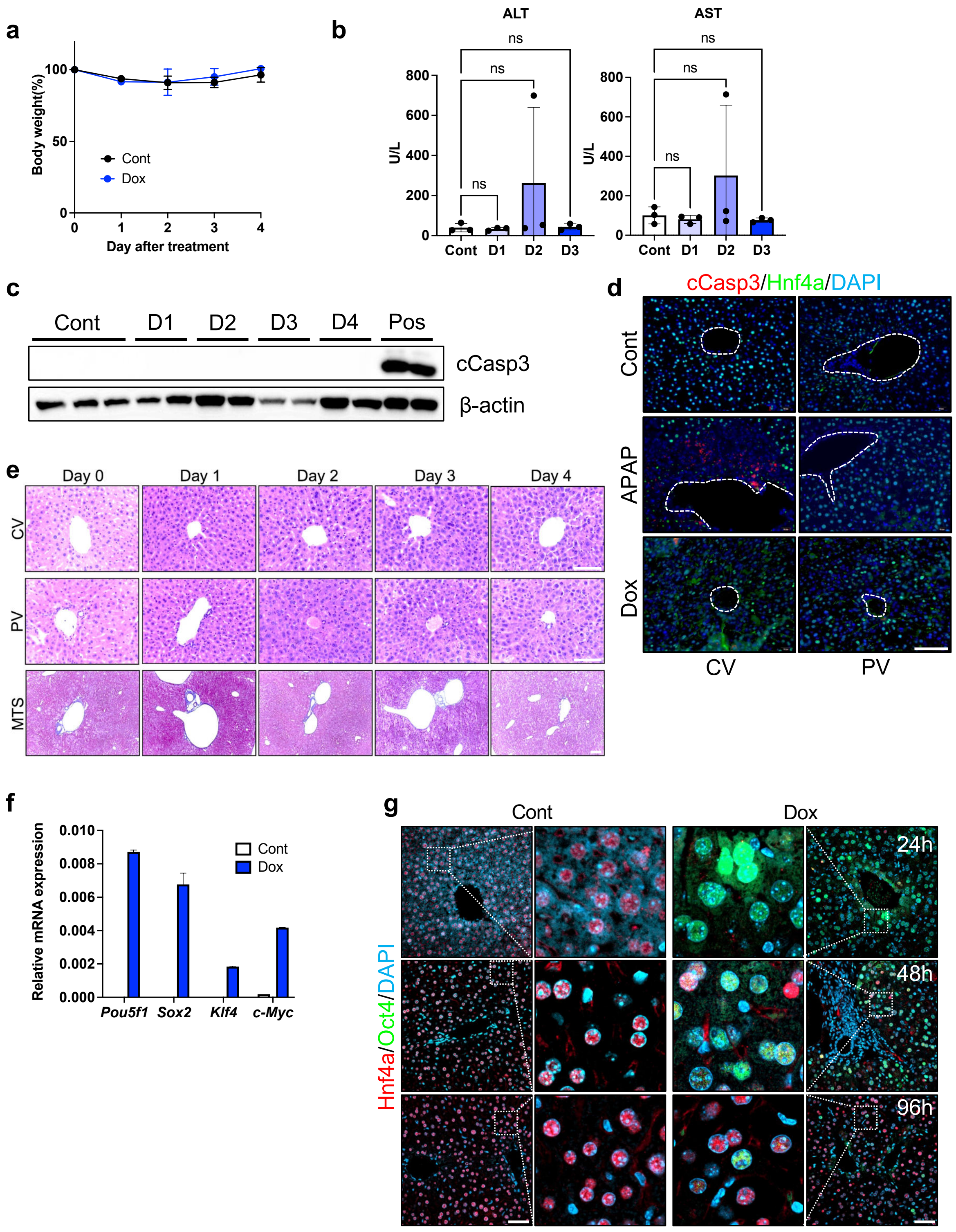


**Supplementary Figure 1. Evaluation of systemic and hepatic responses following short-term OSKM induction in mouse liver.** (a) Body weight analysis with or without Dox treatment for 4 days (n = 3). (b) Biochemical assays for serum ALT and AST levels (n = 3)

after Dox treatment for 3 days. (c) Immunoblot analysis of cleaved Caspase-3 in mouse liver sampled at indicated timepoint after Dox treatment. Positive control (Pos) stands for injured liver (48 h post 300 mg/kg AILI). (d) Immunofluorescence of cleaved Caspase-3 (Red) and Hnf4A (Green) in mouse liver. DAPI stains the nuclei. Dotted line marks the central vein. Scale bar = 100 μm. (e) H&E staining (Upper) and Masson's trichrome staining (Bottom) of mouse liver sampled at indicated timepoints after Dox treatment. Scale bar = 100 μm. (f) Relative RNA expression of OSKM in mouse liver sampled 3 days after Dox treatment. (g) Immunofluorescence of Hnf4A (Red) and Oct4 (Green) in mouse liver. DAPI stains the nuclei. Mouse livers were sampled at indicated time points. 24h; Doxycycline (0.15 mg/ml in drinking water for 24 hours) treated mice. 48h; Doxycycline (0.15 mg/ml in drinking water for 48 hours) treated mice. 96h; Doxycycline (0.15 mg/ml in drinking water for 72 hours + rest for 24 hours) treated mice. Statistical analysis was performed using one-way ANOVA: ns, not significant.

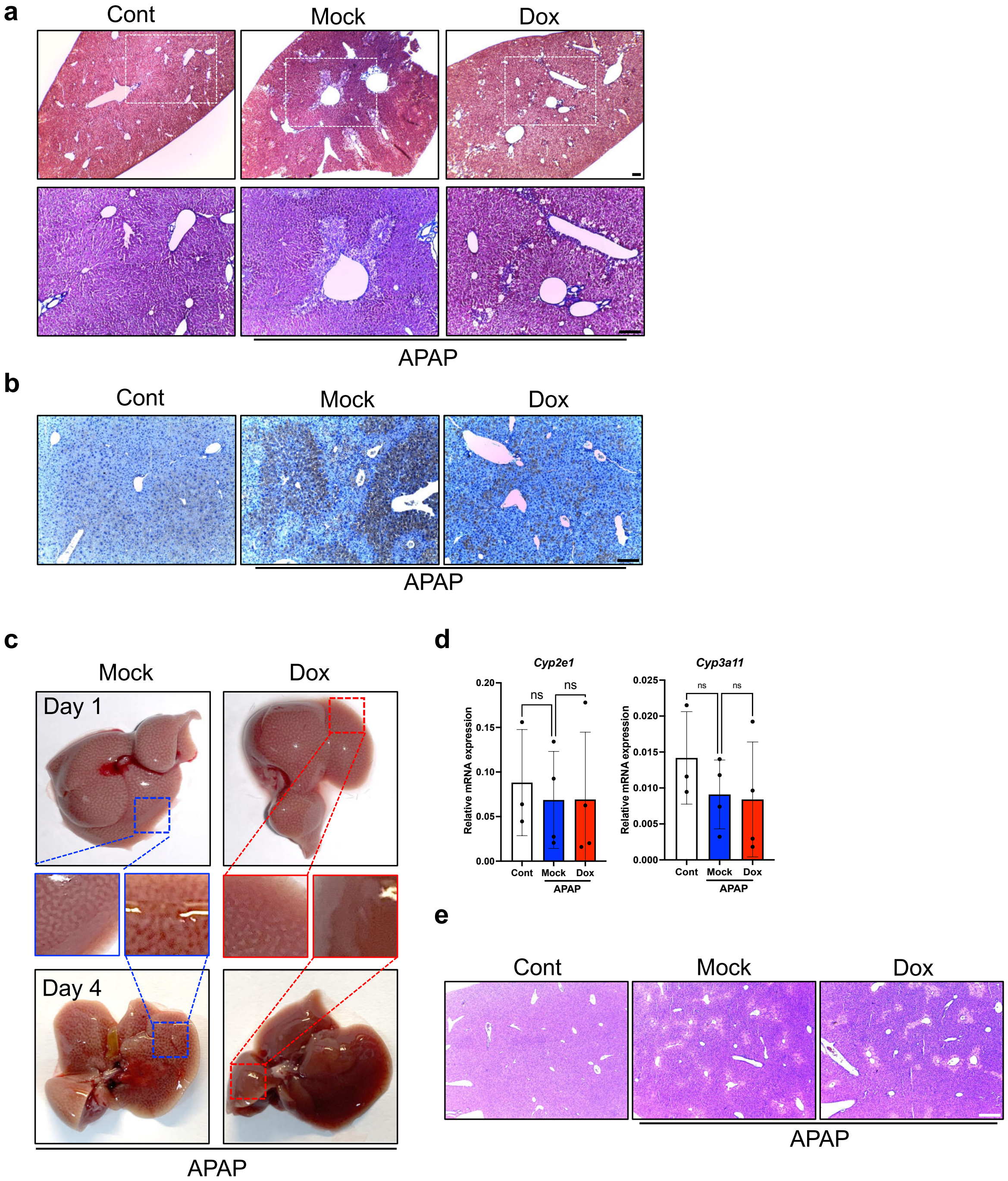


**Supplementary Figure 2. Histological and molecular assessment of reprogramming-mediated modulation in acetaminophen-induced liver injury.** (a) Masson's trichrome staining of mouse liver. 40x and 100x magnification, respectively. (b) Immunohistochemistry of 4-HNE in mouse liver samples collected 48 hours post-AILI. Cont; normal mice, Mock; APAP (300 mg/kg I.P. injection) treated mice, Dox; APAP (300 mg/kg I.P. injection) treated + Doxycycline (0.15 mg/mg in drinking water) treated mice. Scale bar = 100 μm (Control group: n = 3, Mock, Dox group: n = 6). (c) Gross appearance of mouse liver. Insets indicate the distinct lesions induced by AILI. Mock; APAP (500 mg/kg I.P. injection) treated mice, Dox; APAP (500 mg/kg I.P. injection) treated + Doxycycline (0.15 mg/mg in drinking water) treated mice. (d) Relative mRNA expression of *Cyp2e1* and *Cyp3a11* in mouse liver samples collected 6 hours post-AILI. Cont; normal mice, Mock; APAP (300 mg/kg I.P. injection) treated mice, Dox; APAP (300 mg/kg I.P. injection) treated + Doxycycline (0.15 mg/mg in drinking water) treated mice (n = 3 ~ 4). (e) H&E staining of mouse liver samples collected 6 hours post-AILI. Scale bar = 250 μm (n = 3 ~ 4).

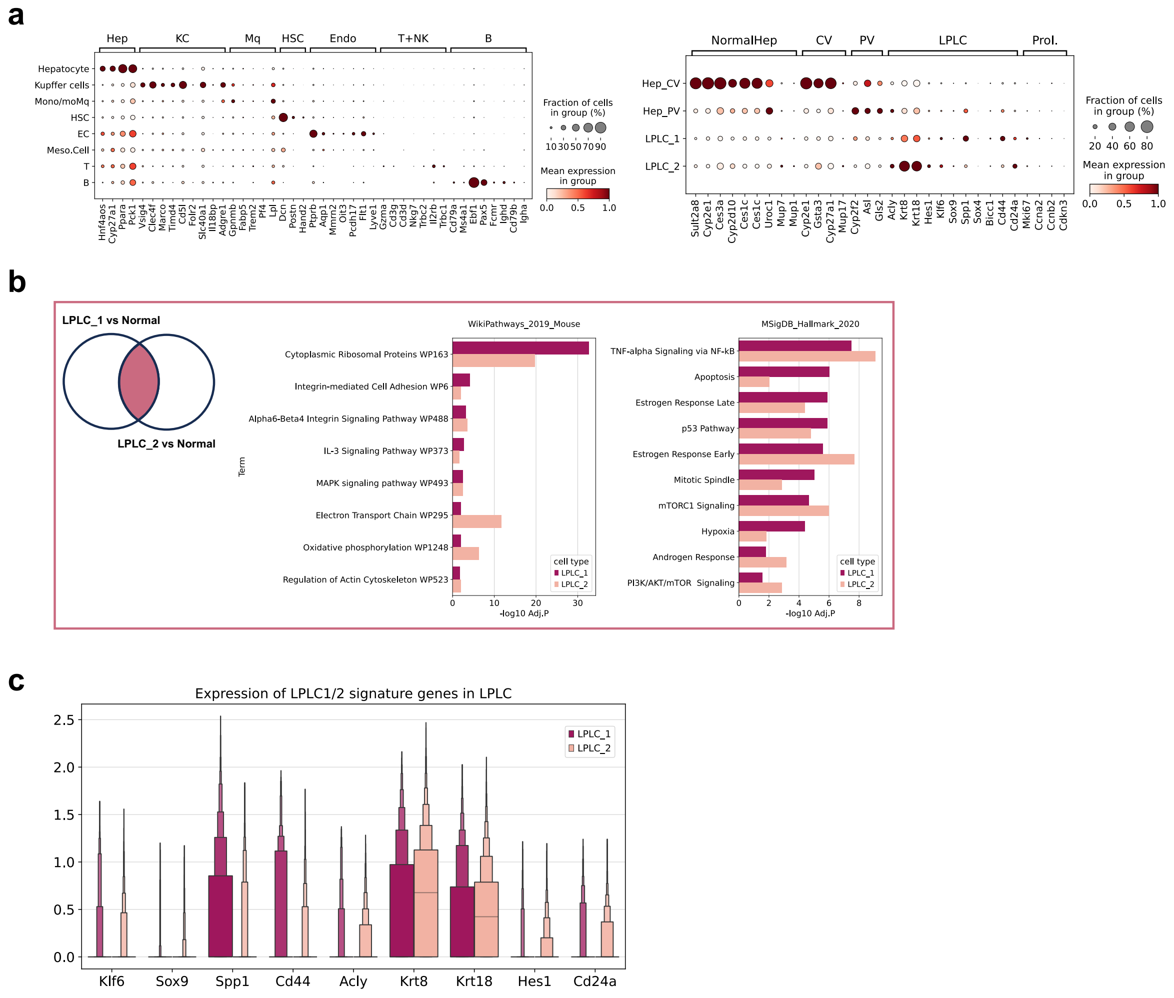


**Supplementary Figure 3. Identification of features of Liver cell types and hepatocyte subpopulations in snRNA-seq data.** (a) Marker genes used for annotating main cell type lineages in Liver (left) and hepatocyte subpopulations (right). (b) Significantly upregulated terms in both LPLC_1 and LPLC_2 were presented. (c) Expression of marker genes for LPLC_1 and LPLC_2 were presented as boxen plot.

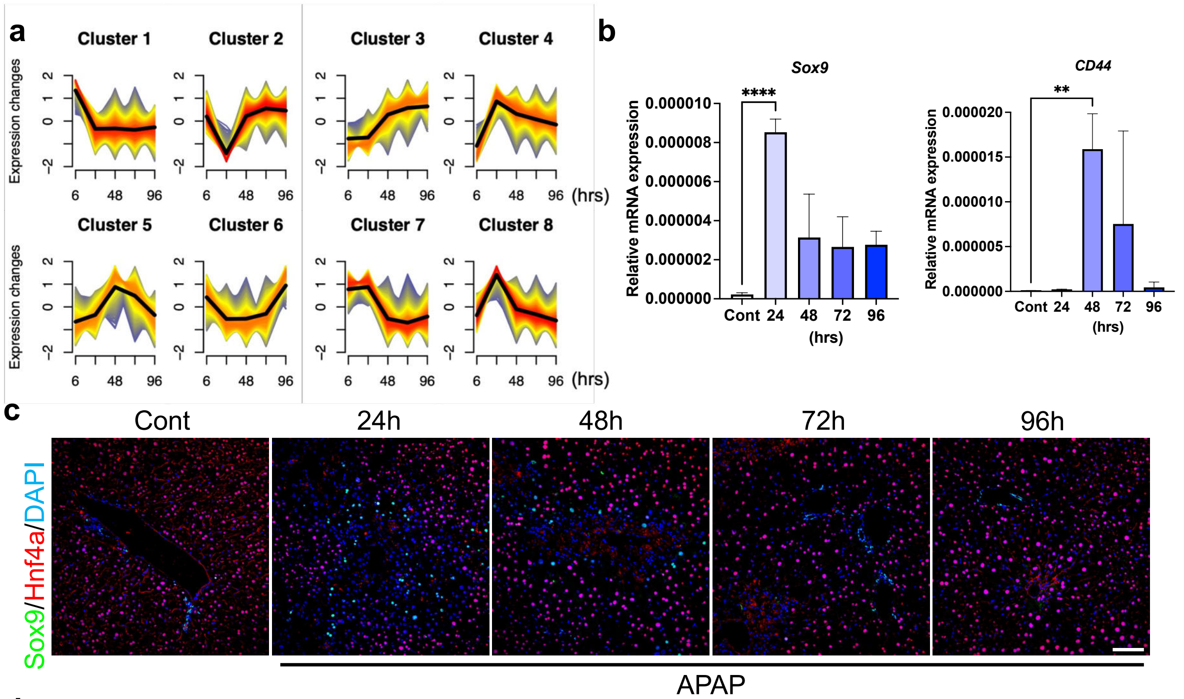


d

| Gene | *Sox9* | *Cd44* | *Ly6a* | *Il6* | *Epcam* | *Cd133* |
|---|---|---|---|---|---|---|
| **Cell type** | LPLCs (Liver progenitor-like cells) | Small hepatocyte | Hepatic Sca-1$^{+}$ progenitor cells | HcPCs (HCC progenitor cells) | Oval cells or Atypical ductal cells(ADCs) | Hepatic progenitor cells |
| **PMID** | 25312494 | 16580085 | 19233130 | 24120137 | 23520387 | 36266314 |

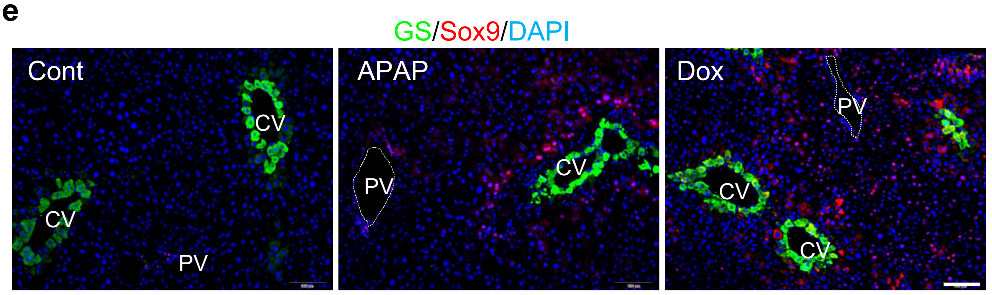


**Supplementary Figure 4. Transcriptomic and histological identification of liver progenitor-like cells induced by partial reprogramming or AILI.** (a) Soft clustering based on gene expression profile post-AILI. DEGs were clustered using 'Mfuzz', threshold was set as 'alpha score > 0.55'. Only genes that have higher than 0.55 alpha score were used on further analysis. (b) Relative mRNA expression of *Sox9* and *Cd44* in whole liver lysate sampled at indicated timepoint post-AILI (n = 3). (c) Immunofluorescence of Hnf4A (Red) and Sox9 (Green) in liver sections sampled at indicated timepoint post-AILI (300 mg/kg I.P. injection). DAPI (Blue) stains the nuclei. Scale bar = 100 μm. (d) List of LPLC markers. 'Cell type' stands for nomenclatures from each reference. (e) Immunofluorescence of Sox9 (Red) and Glutamine synthetase (Green) in liver sections. DAPI (Blue) stains the nuclei. Cont; normal mice, APAP; APAP (48 hours after 300 mg/kg I.P. injection) treated mice, Dox; Doxycycline (0.15 mg/mg in drinking water for 72 hours) treated mice. Scale bar = 100 μm. Statistical analysis was performed using one-way ANOVA: $p < 0.01$(**), $p < 0.0001$(****).

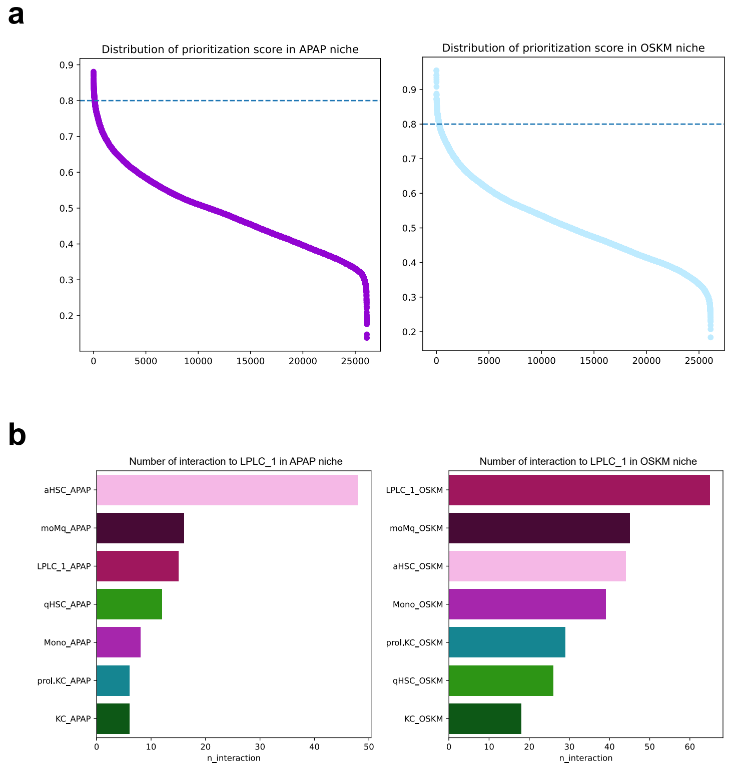


**Supplementary Figure 5. In silico prediction of potential signaling sources for LPLCs.** (a) Prioritization scores for each ligand-receptor pair were presented on scatter plots. In each condition, threshold was set as 0.8. Only ligand-receptor pairs that have higher than 0.8 prioritization score were used on further analysis. (b) Number of interactions to LPLC_1 in each niche. Each cell type indicates signaling sender for LPLC_1.

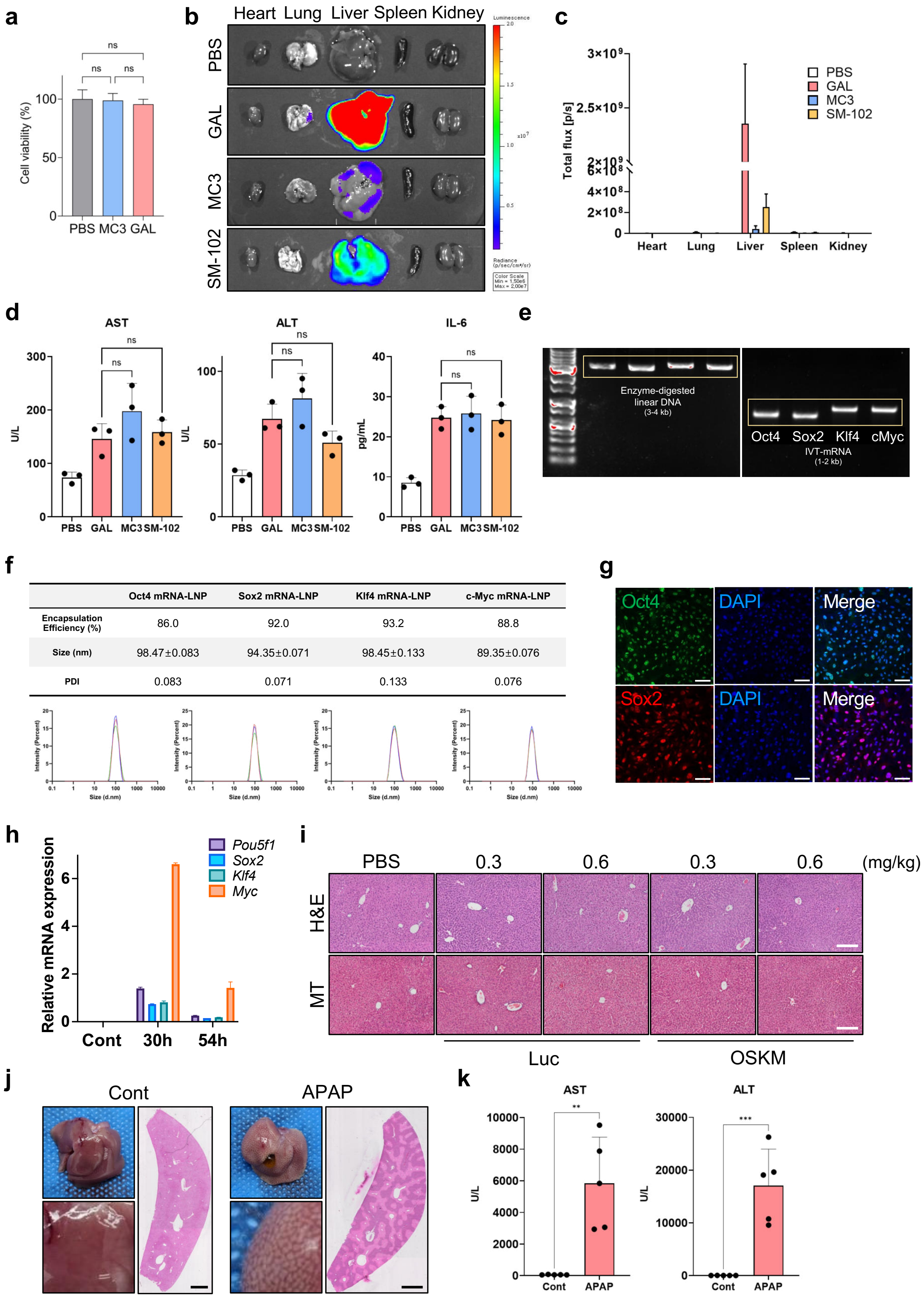


| | Oct4 mRNA-LNP | Sox2 mRNA-LNP | Klf4 mRNA-LNP | c-Myc mRNA-LNP |
|---|---|---|---|---|
| Encapsulation Efficiency (%) | 86.0 | 92.0 | 93.2 | 88.8 |
| Size (nm) | 98.47±0.083 | 94.35±0.071 | 98.45±0.133 | 89.35±0.076 |
| PDI | 0.083 | 0.071 | 0.133 | 0.076 |

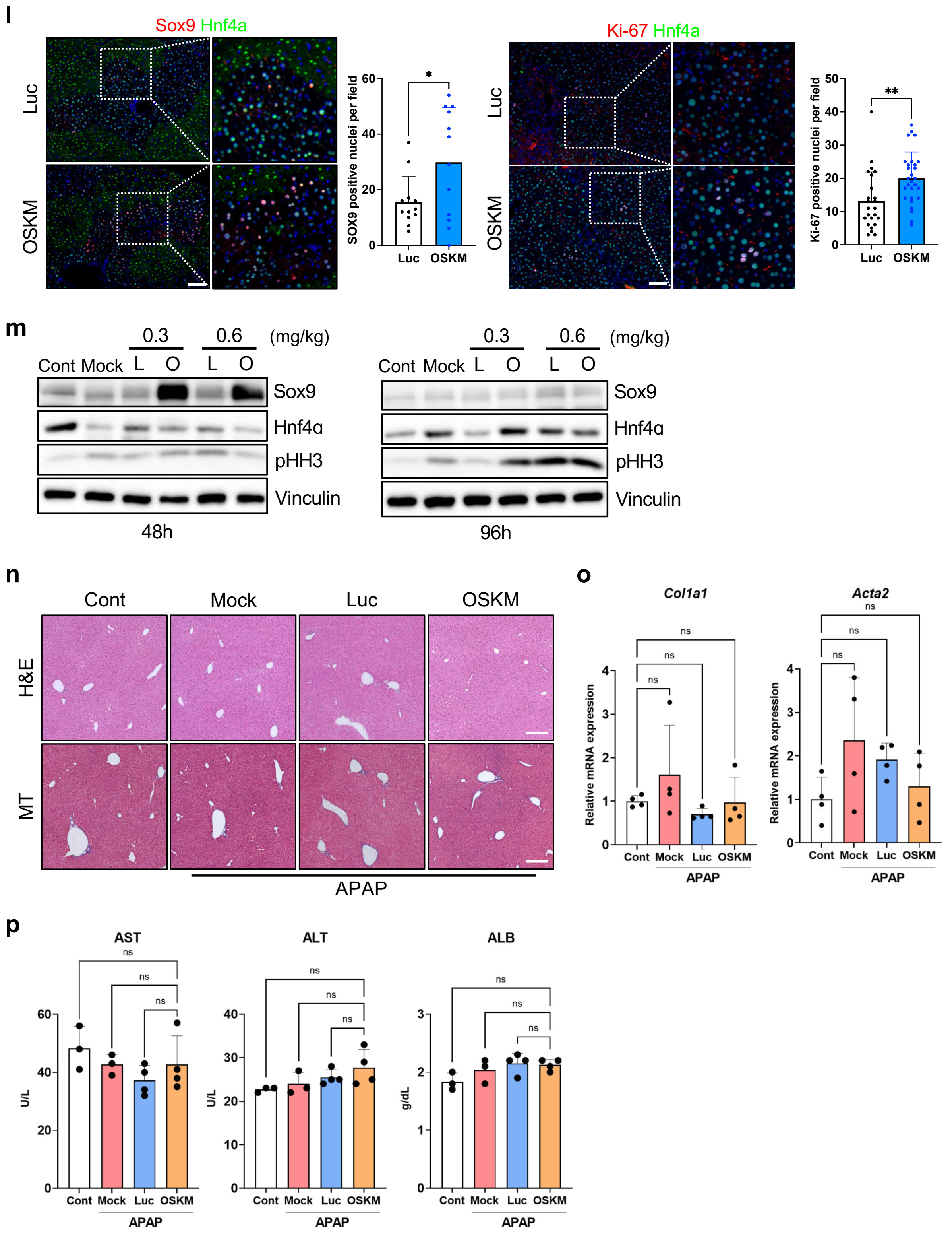


**Supplementary Figure 6. Characterization and validation of OSKM mRNA-LNPs in an acute liver injury mouse model.** (a) MTT assay for measuring viability of MEFs 24 h following treatment with LNPs loaded with firefly luciferase mRNA (1 μg mRNA for 100,000 cells, n = 4). PBS; PBS treatment, MC3; DLin-MC3-DMA LNP treatment, GAL; C12-SPM-

GAL LNP treatment. (b) Representative images of ex vivo organ bioluminescence at 6 hours following intravenous administration of LNPs loaded with firefly luciferase mRNA in C57BL/6 mice (0.3 mg/kg). PBS; PBS-injected mice, GAL; C12-SPM-GAL LNP-injected mice, MC3; DLin-MC3-DMA LNP-injected mice, SM-102; SM-102 LNP-injected mice. (c) Total flux of ex vivo organs collected 6 hours following intravenous administration of LNPs (n = 4). (d) Biochemical assays for serum AST and ALT levels (n = 3) and enzyme-linked immunosorbent assay (ELISA) for serum IL-6 levels (n = 3) at 6 hours post-LNP administration. (e) Agarose gel electrophoresis analysis of NotI restriction enzyme-digested linear OSKM DNA and in vitro transcribed OSKM mRNA. (f) Physicochemical properties of each OSKM mRNA-LNP; encapsulation efficiency (%), particle size (nm), and polydispersity index (PDI) (n = 3, mean ± SD). (g) Immunocytochemistry of Oct4 and Sox2 proteins 24 h after transfection of C12-SPM-GAL LNP loaded with OSKM mRNA into MEFs (1 μg total OSKM mRNA for 100,000 cells, 0.25 μg for each mRNA). DAPI stains the nuclei (scale bars = 100 μm). (h) Quantitative PCR analysis of Oct4 (Pou5f1), Sox2, Klf4, and c-Myc in the liver of C57BL/6 mice 30 and 54 h post-LNP injection (0.3 mg/kg) (n = 5). PBS-treated mice were used as negative control (Cont group). (i) Hematoxylin and Eosin (H&E) staining and Masson's trichrome (MT) staining of liver samples collected 54 h after injection of C12-SPM-GAL LNPs at indicated mRNA doses (scale bars = 200 μm). Luc; Luc mRNA-LNP-injected mice, OSKM; OSKM mRNA-LNP-injected mice. (j) Representative images of the whole liver (left) and H&E staining of whole liver sections (right) one day after APAP administration (400 mg/kg) (scale bars = 1 mm). (k) Blood chemistry analysis of serum AST and ALT levels one day after APAP administration (400 mg/kg) (n = 5). (l) Immunostaining of Sox9 and Ki-67 co-stained with Hnf4α in the liver samples collected 48 h post-APAP administration (30 h post-LNP injection). The graphs show quantification data of Sox9- and Ki-67-positive nuclei per field (n = 3, 4 images per mouse were analyzed) (scale bars = 100 μm). (m) Immunoblot analysis for Sox9, Hnf4α, and pHH3 in the liver collected 48 and 96 h post-APAP administration (30 and 78 h post-LNP injection at indicated mRNA doses). Cont; normal mice, Mock; APAP injury + PBS-injected mice, L; APAP injury + Luc mRNA-LNP-injected mice, O; APAP injury + OSKM mRNA-LNP-injected mice. Vinculin was used as an internal control. (n) Hematoxylin and Eosin (H&E) staining and Masson's trichrome (MT) staining of liver samples collected 30 days post-treatment (scale bars = 200 μm). Cont; normal mice, Mock; APAP injury + PBS-treated mice, Luc; APAP injury + Luc mRNA-LNP-treated mice, OSKM; APAP injury + OSKM mRNA-LNP-treated mice. (o) Quantitative PCR analysis of fibrosisrelated genes in the liver 30 days post-treatment ($n = 4$). (p) Biochemical assays measuring serum AST, ALT, and albumin levels ($n = 3 \sim 4$) in mice 30 days post-treatment. Statistical analyses were performed using one-way ANOVA (Supplementary Fig. 6a, d, o, p) or a Student's t-test (Supplementary Fig. 6k, l): ns, not significant, $p < 0.05$(*), $p < 0.01$(**), $p < 0.001$(***).
